# Structural and functional analysis of lorlatinib analogs reveals roadmap for targeting diverse compound resistance mutations in ALK-positive lung cancer

**DOI:** 10.1101/2021.07.16.452681

**Authors:** Aya Shiba-Ishii, Ted W Johnson, Ibiayi Dagogo-Jack, Mari Mino-Kenudson, Theodore R Johnson, Ping Wei, Scott L Weinrich, Michele A McTigue, Makeba A Walcott, Linh Nguyen-Phuong, Kristin Dionne, Adam Acker, Lesli Kiedrowski, Andrew Do, Jennifer L Peterson, Jaimie L Barth, Beow Y Yeap, Justin F Gainor, Jessica J Lin, Satoshi Yoda, Aaron N Hata

## Abstract

The treatment approach to advanced, ALK-positive non-small cell lung cancer (NSCLC) utilizing sequential ALK tyrosine kinase inhibitors (TKIs) represents a paradigm of precision oncology. Lorlatinib is currently the most advanced, potent and selective ALK tyrosine kinase inhibitor (TKI) in the clinic. However, tumors invariably acquire resistance to lorlatinib, and after sequential ALK TKIs culminating with lorlatinib, diverse refractory compound *ALK* mutations can emerge. Here, we determine the spectrum of lorlatinib-resistant compound *ALK* mutations identified in patients after treatment with lorlatinib, the majority of which involve *ALK* G1202R or I1171N/S/T. By assessing a panel of lorlatinib analogs against compound ALK mutant *in vitro* and *in vivo* models, we identify structurally diverse lorlatinib analogs that harbor differential selective profiles against G1202R- versus I1171N/S/T-based compound *ALK* mutations. Structural analysis revealed that increased potency against compound mutations was achieved primarily through two different mechanisms of improved targeting of either G1202R- or I1171N/S/T-mutant kinases. Based on these results, we propose a classification of heterogenous *ALK* compound mutations designed to focus the development of distinct therapeutic strategies for precision targeting of compound resistance mutations following sequential TKIs.

## INTRODUCTION

Anaplastic lymphoma kinase (*ALK*) gene fusions are the driver oncogene identified across diverse tumor types, including in 3-5% of lung adenocarcinoma (*1–3*). In these tumors, aberrant expression of a constitutively active ALK fusion protein provides a potent oncogenic stimulus that promotes dysregulation and hyperactivation of downstream growth and survival pathways (*1*). Over the past decade, the treatment strategies for metastatic *ALK* fusion-positive (ALK-positive) non-small cell lung cancer (NSCLC) have rapidly evolved with the development of successive generations of increasingly potent and selective ALK tyrosine kinase inhibitors (TKIs) (*4–9*). Multiple randomized studies have demonstrated the superiority of next-generation, highly potent ALK TKIs as initial therapy for metastatic ALK-positive NSCLC (*7–9*). The significant strides made towards optimizing therapeutic targeting of ALK in this short timeframe have resulted in dramatic improvements in patient survival that are unprecedented in metastatic lung cancer (*10–12*) and yielded lessons on targeted therapy applicable across cancers.

Despite marked initial responses to ALK TKIs in most patients, tumors inevitably develop drug resistance leading to disease progression. In 50% to 60% of patients treated with second-generation ALK TKIs (e.g., ceritinib, alectinib, brigatinib), resistance results from the acquisition of secondary mutations in the ALK kinase domain (*13*). These secondary mutations restore ALK activity in the presence of an ALK inhibitor—so-called “on-target” resistance—and the tumors retain dependency on ALK signaling. In particular, the *ALK* G1202R mutation in the solvent front accounts for approximately one-half of on-target resistance across all second-generation ALK TKIs, and I1171T/N/S has been found in 10-15% of patients progressing on alectinib (*13*). Lorlatinib is a third-generation ALK TKI that has demonstrated preclinical potency and clinical efficacy against almost all single *ALK* mutations, including *ALK* G1202R and I1171N/S/T (*13–17*). Of all currently FDA-approved ALK TKIs, lorlatinib is considered the most effective treatment option for ALK-positive NSCLCs refractory to a second-generation ALK TKI (*16*). However, resistance to lorlatinib also emerges. Recent work by our group and others has demonstrated that sequential therapy with a second-generation ALK TKI followed by lorlatinib can drive the development of compound *ALK* resistance mutations (*18–20*). As these tumors are expected to retain dependency on ALK signaling, the development of novel, fourth-generation ALK TKIs that can overcome compound mutation-mediated lorlatinib resistance are urgently needed for patients with ALK-positive NSCLC.

Here, we describe the spectrum of compound *ALK* mutations detected in biopsies from patients with ALK-positive NSCLC treated with lorlatinib. Using Ba/F3 and patient-derived models harboring compound mutant EML4-ALK, we identify lorlatinib analogs with activity against recurrent *ALK* compound mutants involving either G1202R or I1171N/S. These results provide new insights into the clinical and biological significance of compound *ALK* mutations and provide a conceptual framework to guide the development of next-generation inhibitors that can overcome heterogeneous *ALK* compound mutations.

## RESULTS

### Lorlatinib resistance cohort

In our initial description of the molecular landscape of resistance to lorlatinib, approximately one-third of tissue biopsies harbored double or triple *ALK* resistance mutations (*18*). Here, we analyzed a larger updated dataset comprised of 47 patients with lorlatinib-resistant tissue biopsies to determine the prevalence of ALK-positive tumors with ≥2 *ALK* mutations. Demographic characteristics of this group of patients are summarized in **Table S1**.

The majority of patients (85%) received lorlatinib in the third- or later-line setting. All patients had received at least one prior second-generation ALK TKI, and 77% had received prior crizotinib followed by at least one second-generation ALK TKI (**Table S1**). The median time to treatment discontinuation (TTD) of lorlatinib was 8.5 months (95% CI, 6.4-16.1) and the median time to progression (TTP) on lorlatinib was 6.8 months (95% CI, 5.5-10.0), consistent with the known efficacy of lorlatinib in this patient population in the phase 2 trial (*16*). Among 21 patients who had known baseline *ALK* mutation status with a pre-lorlatinib biopsy, both the TTP and TTD were significantly longer in patients with known baseline *ALK* mutations in the tumor (pre-lorlatinib) as compared to those without known baseline *ALK* mutations (TTP: 7.8 months vs 2.8 months, HR 0.363, p<0.0001; TTD: 13.5 months vs 4.1 months, HR 0.128, p=0.0002) (**Fig. S1)**, again concordant with the phase 2 trial results (*17*).

### Landscape of compound *ALK* mutations in lorlatinib-resistant tumors

A total of 48 lorlatinib-resistant tissue biopsies from 47 patients were assessed for resistance mechanisms. Two (4%) demonstrated evidence of histologic transformation: one case from adenocarcinoma to small cell, and another from adenocarcinoma to squamous cell. One case was known to have neuroendocrine histology at initial diagnosis, and this was retained at resistance to lorlatinib. All remaining cases showed non-small cell histology without evidence of histologic transformation.

*ALK* kinase domain mutations were identified in 23 biopsies (48%), of which 14 (29%) were found to harbor double or triple *ALK* resistance mutations (**Fig. 1A-B**, **Table S2**) including the previously unreported *ALK* C1156Y/G1269A (isolated in an ENU mutagenesis screen by our group (*18*) but not previously identified in clinic) and *ALK* G1202R/C1156Y. Using a variety of sequencing techniques as annotated in **Table S2**, we were able to confirm that 9 of the 14 compound mutations were *in cis* (**Fig. 1C**, **Table S2**); for the remainder, it was not feasible to confirm whether the mutations occurred *in cis* versus *in trans* configuration due to the distance between the mutated nucleotides. While no single predominant compound *ALK* mutation was identified, among the 14 cases with ≥2 *ALK* mutations, 8 (57%) harbored *ALK* G1202R—the most common *ALK* resistance mutation after a second-generation ALK TKI—and 3 (21%) harbored *ALK* I1171N—the second most common *ALK* mutation after alectinib (*13*).

**Figure 1.**
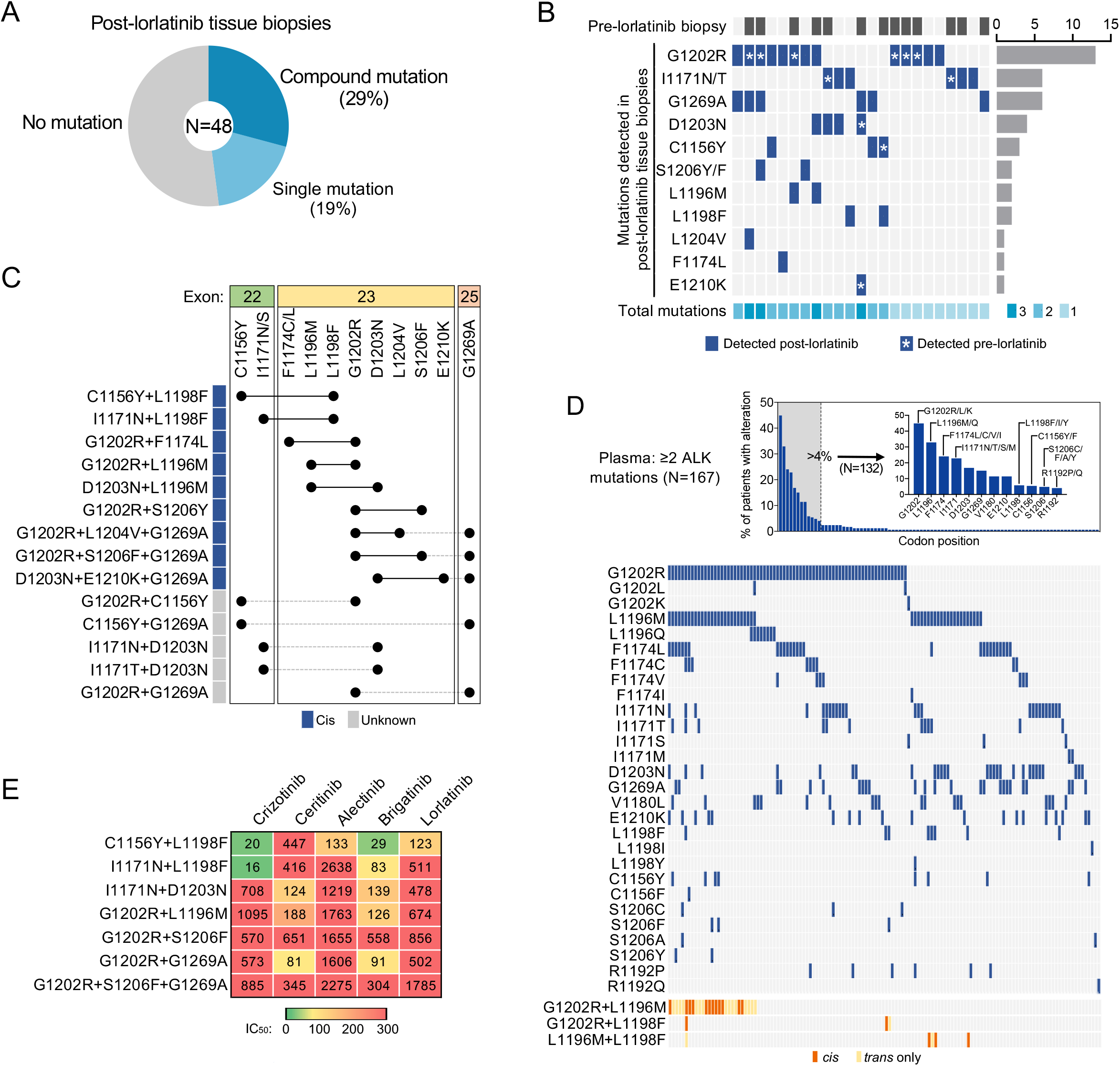
Spectrum of compound *ALK* mutations identified in lorlatinib-resistant biopsies. (A) The frequencies of compound, single, vs no *ALK* mutation detected in lorlatinib-resistant tissue biopsies. (B) Heatmap demonstrating the distribution of *ALK* mutations (dark blue) detected in post-lorlatinib tissue biopsies with at least one *ALK* mutation. Those cases with paired pre-lorlatinib biopsy available are indicated by dark gray (top row). The total number of *ALK* mutations identified are shown by shades of cyan (bottom row). Asterisk indicates the *ALK* mutation was present at baseline (i.e., detected in pre-lorlatinib biopsy). (C) Configuration (*cis* vs *trans*) of *ALK* mutations in tissue biopsies harboring ≥2 *ALK* mutations. (D) *ALK* mutations in an independent cohort of de-identified ALK TKI-resistant plasma samples harboring ≥2 *ALK* mutations. Top panel shows frequency of alterations per codon position. Bottom panel shows co-occurrence of individual mutations at positions altered in greater than 4% of patients. Co-occuring mutations involving closely spaced L1196, L1198 and G1202 positions could be assessed for *cis* (orange) versus *trans* (yellow) configuration. (E) Cellular IC_50_ values of FDA-approved ALK TKIs against clinical compound *ALK* mutations.

The frequency of compound *ALK* mutations was higher in the tumors with acquired vs primary resistance (41% vs 6%, p=0.018). Of the 7 cases with primary lorlatinib resistance (defined as primary progression or disease stabilization for less than 6 months) and paired pre-lorlatinib specimen available, only 1 had harbored a baseline *ALK* resistance mutation, indicative of a pre-existing ALK-independent mechanism in most of these cases (**Table S2**). In contrast, of the 15 cases with acquired lorlatinib resistance and paired pre-lorlatinib tissue, 12 harbored a baseline *ALK* mutation. Of these 12 cases with a baseline *ALK* mutation, an *ALK* mutation was no longer identified in 3 cases at lorlatinib resistance (suggestive of acquired ALK-independent resistance on lorlatinib), and another *ALK* mutation was identified in addition to the pre-existing baseline *ALK* mutation in 5 cases (suggestive of stepwise accumulation of *ALK* mutations on lorlatinib) (**Table S2**, **Fig. 1B**). Additionally, in one patient with an *ALK* I1171T compound mutation for whom a pre-treatment tissue biopsy was not available, the I1171T mutation was detected in plasma prior to initiation of lorlatinib.

### Recurrent compound *ALK* resistance mutations in plasma biopsies

To further assess whether particular *ALK* mutations co-occurred in TKI-resistant ALK-positive NSCLC, we reviewed an independent de-identified Guardant Health dataset comprised of plasma specimens with ≥2 *ALK* mutations. This Guardant cohort included 194 plasma specimens from 167 patients with ALK-positive NSCLC (no treatment history available). The plasma specimens contained between 2 to 10 *ALK* mutations, with the majority (n= 127, 65%) harboring 2 mutations. The most frequent *ALK* mutations detected in plasma overall were: G1202R (44%), L1196M (29%), F1174C/V/L (23%), I1171N/S/T (22%), D1203N (17%), and G1269A (15%) (**Fig. 1D**). *ALK* L1198F was identified in 5% of specimens, of which half also harbored *ALK* G1202R. Because these specimens represent patients treated with different generations of ALK inhibitors, we focused our subsequent analysis on specimens containing *ALK* mutations most commonly associated with alectinib resistance, G1202R (n=73) and I1171N/S/T (n=36). Of the 73 plasma samples with ≥2 *ALK* mutations including G1202R, the *ALK* co-mutations detected in ≥15% of specimens included L1196M (n=27, 37%), F1174C/V/L (n=23, 32%) and I1171N/T (n=15, 21%). Among 36 specimens with ≥2 *ALK* mutations including I1171N/T/S, the *ALK* co-mutations detected in ≥15% of specimens were: G1202R (n=15, 42%), L1196M (n=11, 31%), and V1180L (n=9, 25%), E1210K (n=7, 19%), and D1203N (n=7, 19%). To distinguish whether co-occurring mutations represent compound mutations in the same cell versus independent clones harboring single resistance mutations, we specifically examined the 27 cases with evidence of both G1202R and L1196M, 3 cases with both G1202R and L1198F, and 5 cases with both L1196M and L1198F, all of which lie in close proximity on exon 23 sufficient to be captured on single ctDNA fragments. Collectively, in 17 of 35 (48.6%) cases of co-occurrence, we found evidence that these mutations were *in cis* (12/27 G1202R+L1196M, 2/3 G1202R+L1198F, 3/5 L1196M+L1198F), confirming the recurrent nature of these compound *ALK* mutations in TKI-resistant cases. As the I1171 residue resides on exon 22 whereas the 1180-1210 positions are on exon 23, the allelic relationship between these mutations could not be determined.

### Potency of approved ALK TKIs against compound *ALK* resistance mutations

Our prior work demonstrated that the lorlatinib-resistant *ALK* C1156Y+L1198F double mutation re-sensitized tumor cells to the first-generation inhibitor crizotinib (*21*). In order to assess the sensitivity of additional lorlatinib-resistant compound *ALK* mutations to the currently available, FDA-approved first- and second-generation ALK TKIs, we generated Ba/F3 cell lines harboring distinct *ALK* compound mutations, including the newly discovered G1202R+S1206F+G1269A clinical mutation. Among them, we found distinct patterns of sensitivity vs resistance (**Fig. 1E**, **S2**). Those harboring *ALK* L1198F were sensitive to crizotinib, consistent with prior findings (*21*). On the other hand, compound lorlatinib-resistant *ALK* mutations harboring either G1202R or I1171N without L1198F that were examined here were largely refractory to first- and second-generation ALK inhibitors tested, underscoring the need to discover novel ALK TKIs able to overcome these compound mutations.

### Identification of lorlatinib analogs with activity against compound ALK resistance mutations

To identify novel compounds potent against compound *ALK* mutations, we evaluated a panel of 20 lorlatinib analogs (LAs) (*14, 22*) (**Fig. S3**) in a three-step series of functional screens, using the Ba/F3 models generated above (**Fig. 2A**). In the initial screen, we sought to identify compounds with higher potency than lorlatinib against single *ALK* mutations. This aim was based on our previous observation that the effect of some compound mutations may result from the additive decremental effects of each single mutation on ALK binding and activity (*18*). We then performed a counter screen using PC9 (*EGFR*-mutant NSCLC) and Ba/F3 parental cells to identify compounds with non-ALK-specific cell toxicity. Treatment of Ba/F3 cells harboring single *ALK* mutations with LAs revealed a wide range of effects on cell viability (**Fig. 2B**, **S4**). While all compounds were highly potent against nonmutant EML4-ALK, five of the LAs (LA 1, 14, 16, 17, 18) exhibited IC_50_ values higher than 100 nM in one or more single ALK mutant models. Additionally, three LAs (LA10, 13, 15) reduced viability of Ba/F3 parental cells with IC_50_’s lower than 500 nM with non-ALK-specific cell toxicity (**Fig. 2C**). On the basis of these results, we selected 12 LAs for further testing.

**Figure 2.**
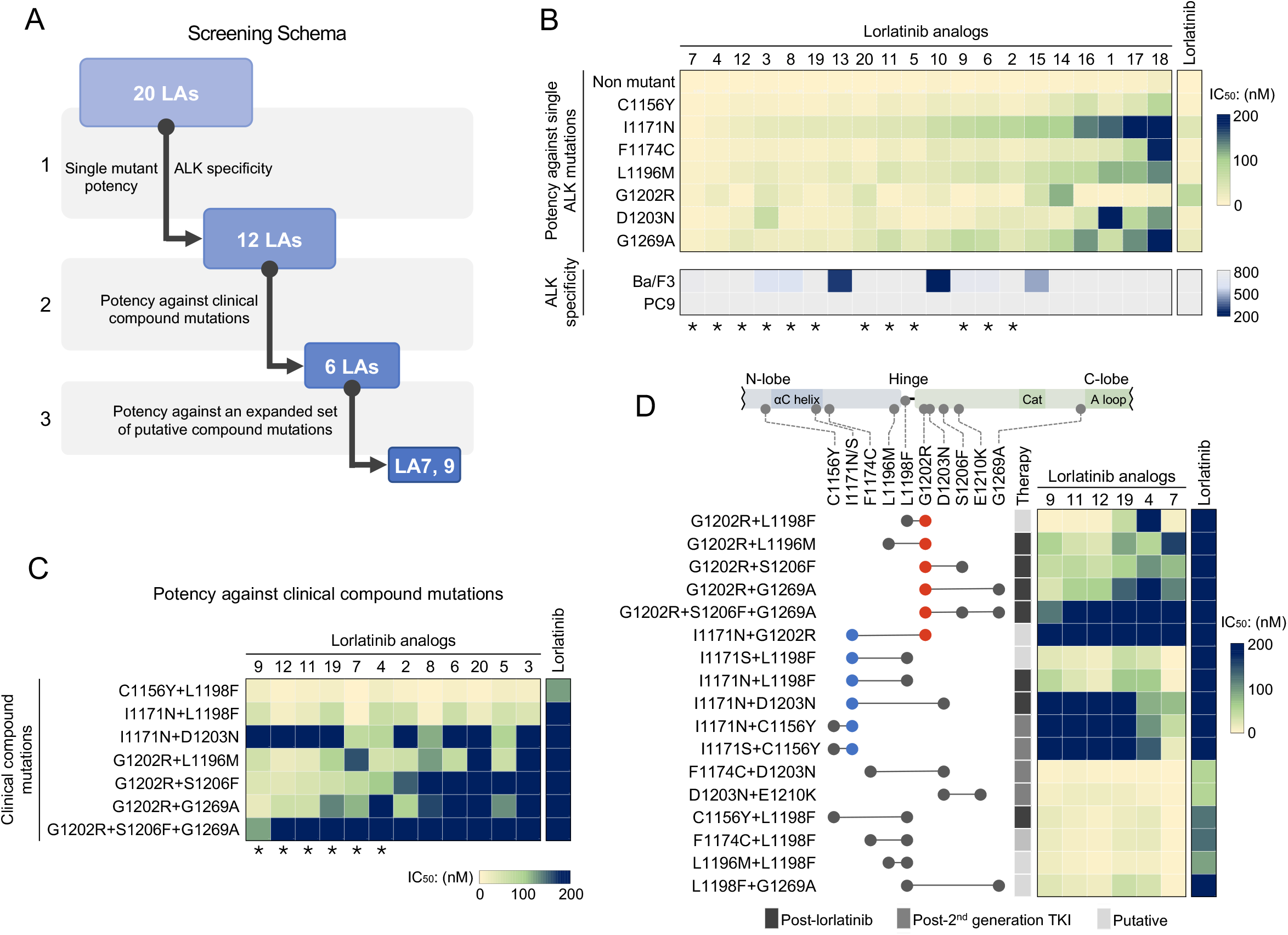
Drug screening of Ba/F3 *ALK* mutation models identifies lorlatinib analogs with increased potency against compound ALK mutations. (A) The schema summarizes the 3-step functional screening of 20 lorlatinib analogs (LAs). (B) Heatmap of cellular IC_50_ values of each LA against single *ALK* mutations (top) and parental Ba/F3 and PC9 cells (bottom). Asterisks indicate the 12 LAs selected for further assessment. (C) Potency of 12 LAs against clinical *ALK* compound mutations. Asterisks indicate the 6 LAs selected for additional validation. (D) Potency of 6 LAs against an expanded set of clinical and putative compound *ALK* mutations. Relative mutation positions within the ALK kinase domain are shown. Red, *ALK* G1202R; blue, *ALK* I1171N/S.

Next, we tested LAs against Ba/F3 models harboring seven distinct compound *ALK* mutations that were identified in post-lorlatinib patients. In general, compound mutation models exhibited higher cellular viability IC_50_ values compared to single mutation models (**Fig. 2C**, **S4**). Among the 12 LAs tested, three (LA9, 11, 12) suppressed cell viability with IC_50_’s lower than 100 nM in five compound mutation models, demonstrating higher potency compared to lorlatinib. Because these three LAs share a similar molecular structure, we additionally selected LA4, 7 and 19 to ensure structural diversity (**Fig. S3**). Using these six candidate LAs, we assessed cell survival of an expanded panel of 17 Ba/F3 cell models harboring clinical and putative *ALK* compound mutations. Similar to our prior results, LAs displayed greater potency with generally lower IC_50_ values against compound mutants than lorlatinib (**Fig. 2D**, **S4**). Among these six LAs, LA7 exhibited the broadest potency, suppressing cell proliferation of 13 compound mutation models with IC_50_’s lower than 100 nM; however, LA7 was not the most potent against all compound mutations and was notably less effective at inhibiting cells harboring G1202R-containing compound mutations. Indeed, by categorizing the sensitivity of compound mutations against LAs, we identified distinct patterns of LA drug efficacy (**Fig. 2D**, **S4**). I1171N/S-containing compound mutations were most sensitive to LA7, whereas LA9/11/12 were most active against G1202R-containing compound mutations. Non-I1171 and G1202 compound mutations—including those containing L1198F—were generally sensitive to all six LAs. Together, these results suggest that distinct compounds may be required to target different categories of *ALK* compound mutations.

### LA7 and LA9 have distinct selectivity profiles against *ALK* compound mutations containing I1171N or G1202R

Based on the Ba/F3 results demonstrating the differential activity of LAs against G1202R and I1171N/S compound mutations, we selected LA7 and LA9 for further validation. Direct comparison of IC_50_ values of LA7 and LA9 (and vs lorlatinib) revealed increased selectivity against I1171N and G1202R single and compound mutants, respectively (**Fig. S6**). When we directly compared the relative potency of LA7 and LA9 against Ba/F3 models classified according to G1202R-based, I1171N/S-based, vs other compound mutants, we confirmed that LA7 suppressed cell viability of I1171N/S and other compound mutation models with greater efficacy, whereas LA9 inhibited G1202R compound mutation models more effectively (**Fig. 3A**), supporting the classification of compound mutations according to the dominant pre-lorlatinib resistance mutation. Next, to investigate the effects of LA7 and LA9 on ALK phosphorylation and downstream signaling pathways, we performed western blotting on G1202R and I1171N compound mutant Ba/F3 models after treatment with LA7 and LA9. While lorlatinib failed to suppress pALK in both compound mutation models, LA7 and LA9 suppressed pALK and downstream pAKT, pERK and pS6 signaling (**Fig. 3B**, **Fig. S7**). Consistent with the cell survival assay results, we again observed a distinct pattern of differential LA activity. LA7 suppressed pALK in I1171N+D1203N and I1171N+L1198F models more effectively than LA9, and conversely, LA9 more potently suppressed G1202R+L1196M, G1202R+G1269A and G1202R+S1206F+G1269A models. Finally, we compared the activity of LA7 and LA9 to clinically available ALK inhibitors. Overall, LA7 and LA9 were the most potent against the seven compound mutations that we identified in post-lorlatinib patients (**Fig. S8**).

**Figure 3.**
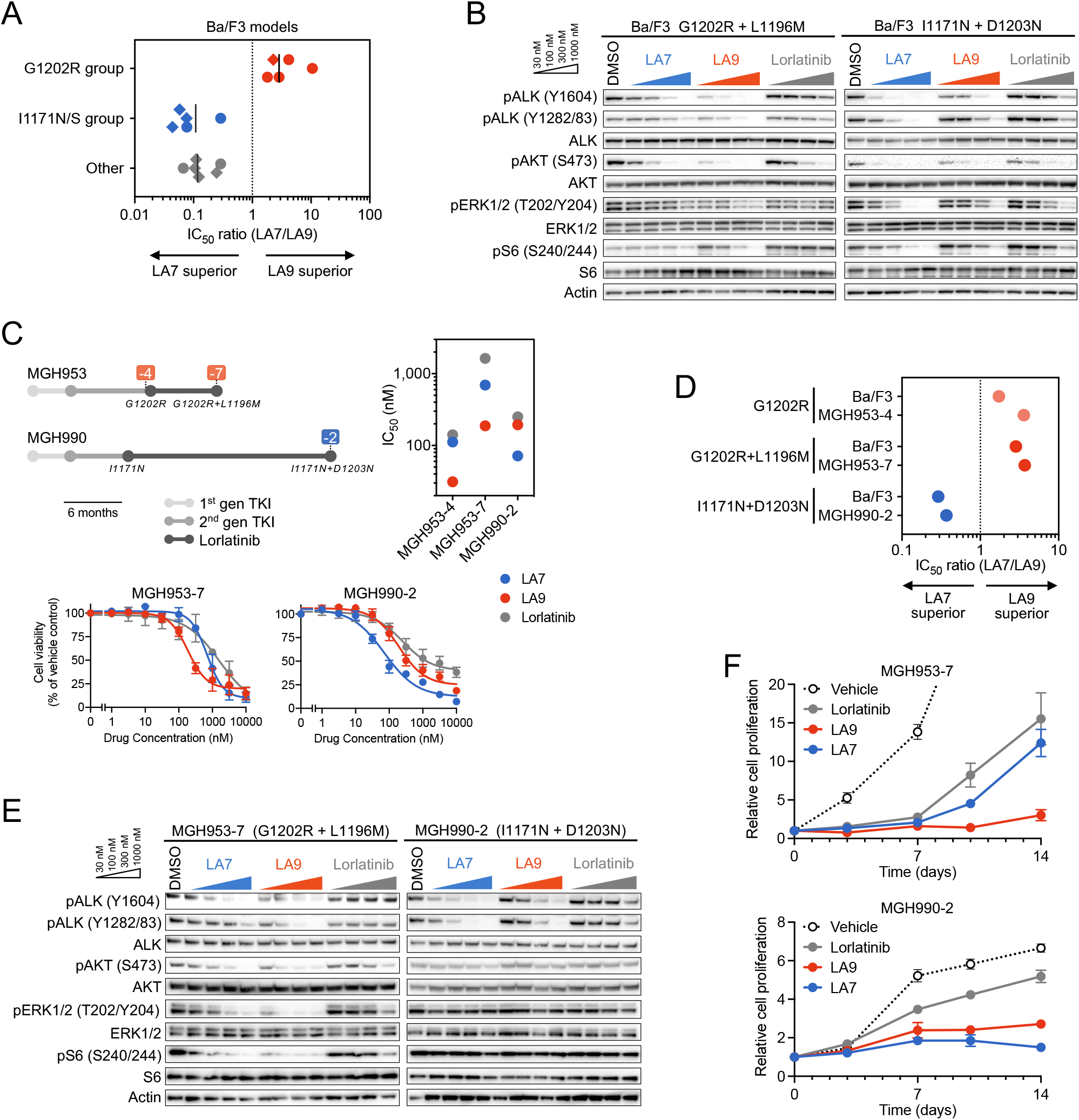
LA7 and LA9 are differentially active against I1171N- and G1202R-containing compound mutations. (A) Selectivity of LA7 and LA9 against Ba/F3 models harboring G1202R (G1202R+L1198F, G1202R+L1196M, G1202R+S1206F, G1202R+G1269A, G1202R+S1206F+G1269A), I1171N/S (I1171S+L1198F, I1171N+L1198F, I1171N+D1203N, I1171N+C1156Y, I1171S+C1156Y), or other compound *ALK* mutations (F1174C+D1203N, D1203N+E1210K, C1156Y+L1198F, F1174C+L1198F, L1196M+L1198F, L1198F+G1269A) calculated by the ratio IC_50_(LA7)/IC_50_(LA9). Diamonds indicate clinical mutations, and circles indicate putative mutations. (B) Western blot analysis performed in Ba/F3 cells expressing ALK G1202R+L1196M or ALK I1171N+D1203N after treatment with DMSO, LA7, LA9, or lorlatinib for 6 hours. (C) MGH953-7 (G1202R+L1196M) and MGH990-2 (I1171N+D1203N) patient-derived cell lines treated with the indicated drugs for 72 hours, with viability measured using CellTiter-Glo assay. Error bars indicate SE. The treatment histories of the MGH953 and MGH990 patients are summarized (top left), and cellular IC_50_ values in MGH953-7, MGH990-2, and MGH953-4 (G1202R) are shown (top right). (D) Selectivity of LA7 and LA9 against patient-derived and Ba/F3 models harboring I1171N vs G1202R mutations, respectively. (E) Western blot analysis performed in MGH990-2 and MGH953-7 cells after treatment with indicated drugs for 6 hours. (F) Cell growth of MGH953-7 and MGH990-2 cells treated with the indicated drugs for 14 days. 300 nM of each drug was used for MGH953-7 and 100 nM of each drug was used for MGH990-2. Error bars indicate SE.

Next, we compared the efficacy of LA7 and LA9 against patient-derived cell lines generated from tumors harboring G1202R single mutation (MGH953-4), G1202R+L1196M compound mutation (MGH953-7) and I1171N+D1203N compound mutation (MGH990-2). Concordant with the results in Ba/F3 models, LA9 inhibited the growth of MGH953-7 harboring G1202R+L1196M most effectively, whereas LA7 was most potent against MGH990-2 harboring I1171N+D1203N (**Fig. 3C**, **S9**). Direct comparison of the selectivity profiles of LA7 and LA9 in the patient-derived cells showed close concordance with the Ba/F3 models harboring the same mutations (**Fig. 3D**). Consistent with the cell viability results, LA7 and LA9 more effectively suppressed pALK signaling in MGH990-2 and MGH953-7 cells, respectively (**Fig. 3E**). To test the ability for LA7 and LA9 to suppress the growth of ALK compound mutants over time, we treated MGH953-7 and MGH990-2 cells with concentrations of LA7, LA9 or lorlatinib predicted to discriminate between compound mutations for 14 days and monitored the effects on cell growth. Consistent with the signaling and short-term viability results, LA9 suppressed cell growth of MGH953-7 (G1202R+L1196M) to a greater extent than LA7 or lorlatinib (**Fig. 3F**). Conversely, LA7 inhibited cell growth of MGH990-2 (I1171N+D1203N) more effectively than LA9 or lorlatinib.

Finally, we investigated whether LA7 and LA9 are able to inhibit the growth of ALK-positive NSCLC tumors harboring compound mutations *in vivo*. As expected, MGH953-7 primary patient-derived xenograft (PDX) tumors harboring G1202R+L1196M were resistant to clinically relevant doses of lorlatinib (**Fig. 4A**). To account for differences in plasma protein binding, we estimated the unbound plasma concentrations of lorlatinib, LA7 and LA9 to select comparable doses using a once-daily (QD) dosing regimen (**Fig. S10A, B**). At doses that were well tolerated and confirmed to yield approximately similar unbound systemic exposures as lorlatinib (corresponding to LA7 20 mg/kg, LA9 40 mg/kg; **Fig. S10C, D** **and** **S11**), both LA7 and LA9 initially slowed tumor growth; however, upon extended treatment, LA7-treated tumors regrew, while the LA9-treated tumors continued to be largely suppressed (p=0.0076 for tumor size difference on day 32; **Fig. 4A**). Consistent with these results, immunohistochemical analysis of MGH953-7 xenograft tumors demonstrated that ALK phosphorylation was effectively inhibited in LA9-treated tumors, but not lorlatinib or LA7-treated tumors (**Fig. 4B**, **S12**). Despite multiple attempts, we were unable to establish xenograft tumors from the MGH990-2 (I1171N+D1203N) cell line; therefore, in order to test the efficacy of LA7 *in vivo*, we generated NIH3T3 cells expressing ALK I1171N+D1203N. Consistent with the I1171N+D1203N Ba/F3 and MGH990-2 cell lines, we confirmed that LA7 more potently suppressed ALK phosphorylation compared to LA9 or lorlatinib *in vitro* (**Fig. 4C**). Finally, we established 3T3 I1171N+D1203N xenograft tumors in immunocompromised mice and treated with lorlatinib, LA7 or LA9 using the same doses as those used for the MGH953-7 (G1202R+L1196M) xenograft tumors. LA7 achieved significantly greater tumor growth inhibition in 3T3 I1171N+D1203N tumors compared to LA9 or lorlatinib (**Fig. 4D**).

**Figure 4.**
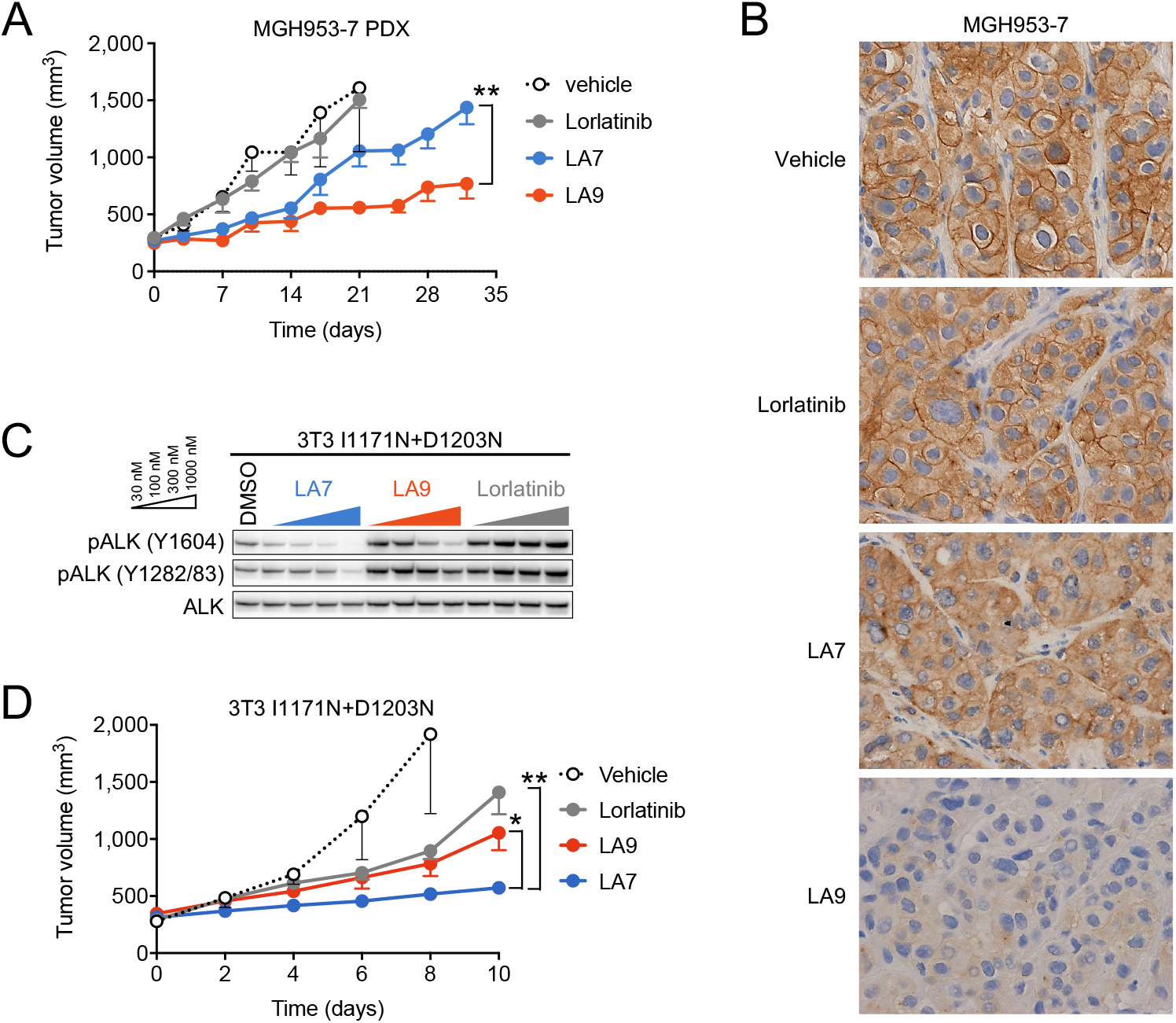
LA7 and LA9 inhibit tumor growth in xenograft tumors bearing *ALK* compound mutations. (A) Change in tumor volume of MGH953-7 (G1202R+L1196M) PDX tumors. Mice were treated with vehicle (*n* = 6), lorlatinib 6 mpk QD (*n* = 5), LA7 20 mpk QD (*n* = 6), or LA9 40 mpk QD (*n* = 6), 5 days per week. Tumor sizes on day 32 were significantly different between LA7 and LA9 (p=0.0076, two-tailed t test). Error bars indicate SE. (B) Immunohistochemistry for pALK performed on MGH953-7 xenograft tumors harvested after 3-day treatment with the indicated drugs. (C) Western blot analysis of NIH3T3 cells expressing ALK I1171N+D1203N and treated with the indicated drugs. (D) Change in tumor volume of NIH3T3 I1171N+D1203N xenograft tumors. Mice were treated with lorlatinib 6 mpk QD, LA7 20 mpk QD, or LA9 40 mpk QD (*n* = 6 mice per group). Tumors treated with LA7 were significantly smaller on day 10 compared to those treated with LA9 (p=0.0133) or lorlatinib (p=0.0018, two-tailed t test). Error bars indicate SE.

### Structural basis for improved potency of LA7 and LA9 against compound mutants

To better understand the mechanistic basis for the selectivity of LA9 and LA7 against G1202R- and I1171N/S/T compound mutants, respectively, we first compared the cellular IC_50_ values of lorlatinib for nonmutant, single and compound mutant ALK. Consistent with previous reports (*13*), both single mutations G1202R and I1171N resulted in relative increases in lorlatinib IC_50_ values (~88-fold and ~43-fold, respectively) compared to nonmutant ALK; however, the absolute IC_50_ values remained within the range of clinical exposures (~200 nM) (*15*) (**Fig. 5A**, **S13A**). Compound mutations resulted in a further relative increase in lorlatinib IC_50_ of approximately 10-fold (range, ~7-24 fold), increasing the absolute IC_50_ values above the upper range of plasma concentrations that are clinically achievable. By comparison, LA7 and LA9 exhibited IC_50_ values against I1171N- and G1202R-containing compound mutations that ranged from ~7 to 138-fold lower than lorlatinib, respectively (**Fig. S13B**). Notably, LA7 and LA9 exhibited a similar relative improvement in potency against I1171N and G1202R single mutations, respectively (~11 and ~18 fold, respectively), suggesting that the improved drug efficacy against compound mutations is predominantly attributable to improved ability to overcome either the I1171N or G1202R single site substitutions. Consistent with this notion, neither drug was capable of overcoming the putative *ALK* I1171N+G1202R compound mutation (**Fig. 2D**).

**Fig. 5.**
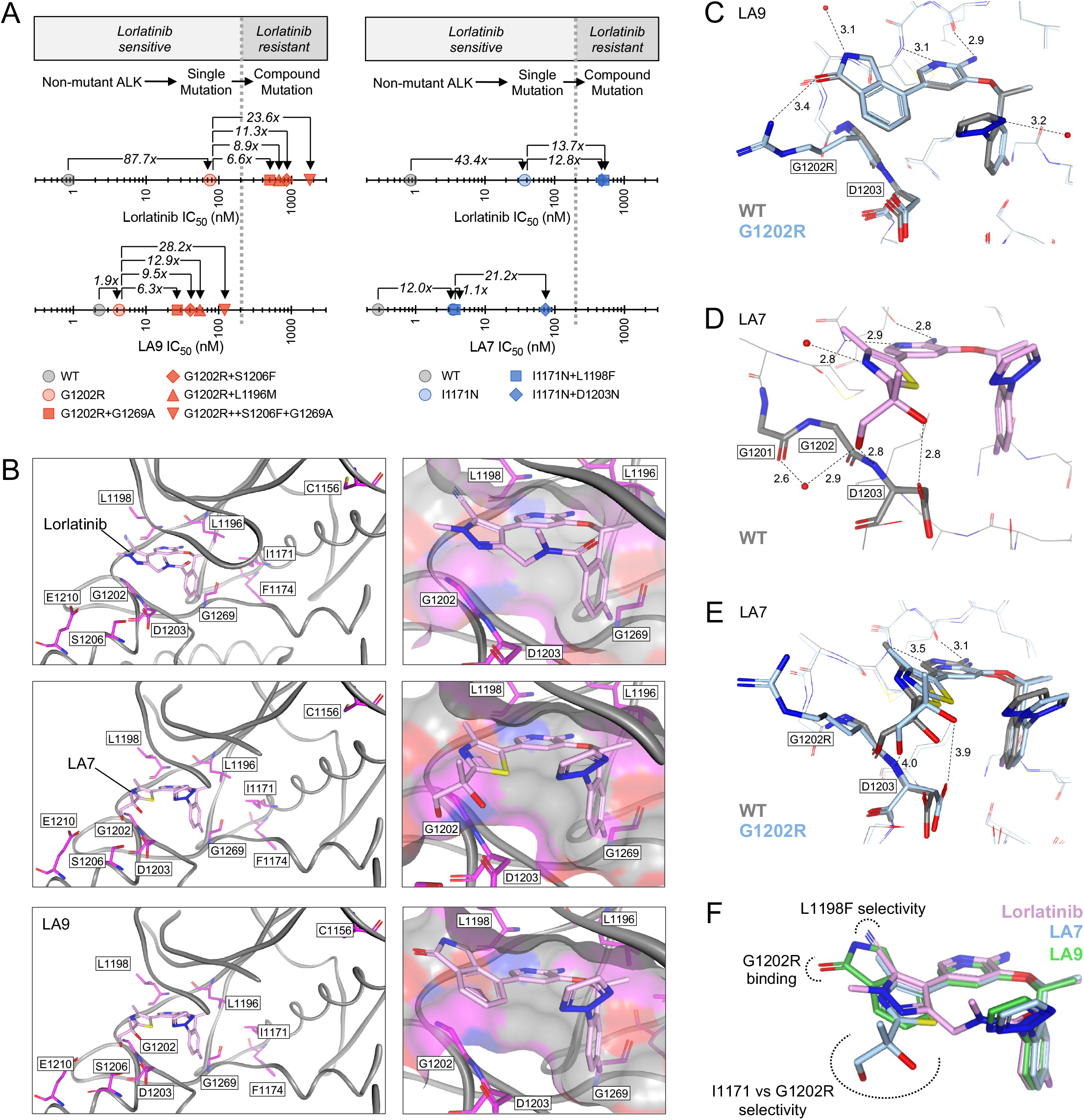
Structural basis for selectivity of LA7 and LA9 against *ALK* compound mutations. (A) Comparison of absolute and fold shift in cellular IC_50_ values for nonmutant, single and compound ALK mutants (corresponding to data shown in Figure S5). G1202R and I1171N single point mutations result in large shift in IC_50_ for lorlatinib; however, this remains within range of clinical exposure (<200nM, indicated by dashed line). Compound mutations result in comparable fold shift in IC_50_ values of lorlatinib, LA7 and LA9 relative to single mutations; however, the improved potency of LA7 and LA9 against single mutations results in substantial improvement in IC_50_ against compound mutations. (B) Co-crystal structures of lorlatinib, LA7 and LA9 in the wild-type ALK kinase domain. Left panels highlight residues mutated in resistant patients. Right panels show structure of the ligand binding pocket. (C) Co-crystal structure of LA9 bound to wild-type ALK (ligand and protein colored gray) superimposed with energy-minimized model of G1202R (ligand and protein colored blue). Solvent front G1202/G1202R and D1203 residues are bolded for emphasis. Productive interactions between both LA9/WT and LA9/G1202R are shown by dotted lines, including a novel interaction between G1202R side chain –NH2 and the lactam carbonyl of LA9 (distances are reported in Angstroms). Note the minimal movement of both ligand (LA9) and protein (G1202R) relative to LA9/WT co-crystal structure. (D) Co-crystal structure of LA7 in WT ALK highlighting productive interactions, including a novel network of hydrogen bonds between thiazole ring hydroxyl groups and solvent front residues (1201-1203, bolded; red spheres represent structural water molecules). (E) Co-crystal structure of LA7 in WT ALK (ligand and protein colored gray) superimposed with energy-minimized model of G1202R (ligand and protein colored blue) showing significant movement of both ligand and protein relative to LA7/WT co-crystal structure. This movement disrupts key protein ligand interactions, including the two hydroxyl groups with G1201 and G1203 shown in panel D. Distances of LA7/G1202R interactions are shown. (F) Overlap of binding modes of lorlatinib, LA7 and LA9 (in WT ALK) revealing orientation of key structural features conferring mutant selectivity.

Next, we examined the structural basis for the improved potency of LA9 and LA7 against G1202R- and I1171N-compound mutants, respectively. Comparison of the co-crystal structures of lorlatinib, LA7 and LA9 bound to nonmutant ALK confirmed similar binding modes of all three ligands (**Fig. 5B**). Previously, we reported that the introduction of the G1202R solvent front mutation increases steric bulk near the pyrazole ring of lorlatinib, leading to partial destabilization of ligand binding and an altered P-loop conformation (*18*). Modeling the G1202R mutation based on the LA9/ALK co-crystal structure revealed that the orientation of the unsubstituted phenyl ring near G1202 enables accommodation of the G1202R substitution without significant steric clash or disruption of the ligand position (**Fig. 5C**). Furthermore, the G1202R side chain amino group makes a new productive interaction with the lactam carbonyl of LA9. In contrast to the predominantly steric constraints introduced by the solvent front G1202R substitution, I1171N lies within the hydrophobic core and confers ALK TKI resistance by increasing the catalytic activity of ALK (*23–25*). Interestingly, LA7 exhibited improved potency against nonmutant ALK as well as I1171N (**Fig. S13**), suggesting favorable interactions between the ligand and wild-type residues. Examination of the co-crystal structure of LA7 bound to nonmutant ALK revealed stabilizing interactions between the two hydroxyl groups of the ligand thiazole ring and the side chain carboxylate of D1203, as well as a hydrogen bond network involving the backbone carbonyl of G1201, the amino nitrogen of D1203 and a structural water (**Fig. 5D**), that are not present with LA9 or lorlatinib. While not involving interactions with G1202 directly, the substitutions on the thiazole ring of LA7 are positioned adjacent to the solvent front residues and result in a significant steric clash upon G1202R substitution. This disrupts key protein-ligand interactions (including hinge binding and the two hydroxyl groups with G1201 and D1203) and causes significant movement of ligand relative to the LA7/ALK co-crystal structure (**Fig. 5E**). Thus, mutually exclusive molecular features of LA7 and LA9 enable improved targeting of I1171N/S/T- and G1202R-class mutations, respectively.

Finally, we examined the structural basis for overcoming L1198F-containing compound mutations. Originally, the lorlatinib nitrile group was installed to improve ALK-selectivity based on the observation that most kinases contain either a phenylalanine or tyrosine corresponding to the position at L1198F (*14*). The L1198F mutation coupled with the rigid nature of the lorlatinib macrocycle causes a steric clash between the phenyl group and nitrile group of the inhibitor, reducing affinity (*21*). Five out of the six selected LAs (**Fig. 2D**), including LA7 and LA9, lack the nitrile and consistently exhibited high potency against L1198F compound mutations (**Fig. S4C**, **S5**). Consistent with this result, structural modeling of L1198F on the LA7 and LA9/ALK co-crystal structures revealed that the mutation had minimal impact on ligand binding (**Fig. S14A-C**). LA4, on the other hand, contains a nitrile but is otherwise identical to LA7 (**Fig. S3**). Comparison of cellular IC_50_ values of LA4 vs LA7 across compound mutants revealed a significant decrease in potency of LA4 compared to LA7 specifically against L1198F-containing compound mutations (**Fig. S14D**). Modeling of L1198F on the LA4/ALK co-crystal structure revealed orientation of the nitrile toward L1198F (**Fig. S14E**), albeit with less steric clash compared to lorlatinib (owing to the more flexible LA4 structure), consistent with the decreased activity against L1198F mutants compared to LA7 (**Fig. S14F**). Collectively, these findings reveal the structural basis for a rational approach to targeting diverse compound resistance mutations classified according G1202R and I1171N (**Fig. 5F**).

## DISCUSSION

Currently, most patients with advanced ALK-positive lung cancer are treated using a sequential therapy paradigm with successive generations of ALK TKIs, often culminating with the third-generation and most potent TKI, lorlatinib. While this sequential targeted therapy approach has enabled prolonged survival and quality of life, acquired resistance to lorlatinib is inevitable, and treatment options following lorlatinib remain an area of critical unmet need. Our current study extends prior work by our group and others demonstrating that in a subset of these patients, acquired resistance is caused by recurrent lorlatinib-refractory *ALK* compound mutations that sequentially evolve from *ALK* single mutant clones arising after prior treatment with first- or second-generation ALK TKIs (*18–20*). Efforts to develop fourth-generation TKIs that can overcome *ALK* compound mutations are underway (*26, 27*); however, the diverse spectrum of *ALK* compound mutations observed in patients after lorlatinib suggests that one approach is unlikely to be effective for all patients. Indeed, our prior finding that the acquisition of a compound *ALK* mutation containing L1198F confers resensitization to crizotinib (*21*) is not always recapitulated with other compound mutations, many of which remain largely refractory to currently available ALK TKIs. By analyzing a structurally diverse set of lorlatinib analogs, we identified molecules with distinct sensitivity patterns against classes of *ALK* compound mutations grouped according to the predominant resistance mutation that exists pre-lorlatinib, G1202R or I1171N/S/T. As these represent the two most common resistance mutations observed in patients progressing on alectinib (the preferred first-line ALK TKI in the recent years), our findings offer a roadmap for rational development of ALK inhibitors targeting subgroups of ALK-positive lung cancers with resistance to lorlatinib.

Comparison of potencies of lorlatinib and lorlatinib analogs against diverse compound mutations, integrated with our analysis of co-crystal structures of lorlatinib analogs bound to ALK, reveals the structural basis for overcoming *ALK* compound mutations. While single residue substitutions in the kinase domain are sufficient to cause resistance to first- and second-generation ALK TKIs, lorlatinib binds more potently and is able to inhibit G1202R and I1171N/S/T resistance mutations at clinically achievable drug exposures (~200 nM) (*15*), However, the relative potency of lorlatinib against these mutations is reduced compared to nonmutant ALK, and our analysis reveals this to be a major factor contributing to clinical resistance conferred by subsequent compound mutations. In virtually all cases, the introduction of a second mutation in ALK that already harbors a G1202R or I1171N single point mutation leads to a further ~10-fold decrease in the potency of both lorlatinib and the lorlatinib analogs. For lorlatinib, this decreased potency translates to a requirement for drug concentrations that are not clinically achievable. On the other hand, for the lorlatinib analogs tested, we identified multiple compounds including LA9 and LA7 with markedly improved potency (≥10-fold) against *ALK* G1202R or I1171N/S/T single mutations. Thus, even with a similar relative reduction in potency incurred upon the introduction of a second mutation, LA9 and LA7 exhibited >10-fold increased potency relative to lorlatinib against diverse G1202R or I1171N/S/T compound mutations, respectively, leading to improved activity *in vitro* and *in vivo*.

From a structural perspective, LA7 and LA9 achieve improved targeting of I1171N/S/T and G1202R substitutions by distinct mechanisms. Kinase domain substitutions may contribute to resistance in complex ways, including by reducing the effectiveness of the ligand’s ability to bind and block ATP, or alternatively by increasing the catalytic efficiency of the kinase (which in turn may derive from changes in substrate binding, ATP binding, or conformational changes in the kinase that favor the active form). Our prior work revealed that the G1202R solvent front mutation in ALK increases steric bulk near the pyrazole ring of lorlatinib, leading to destabilization of ligand binding and an altered P-loop conformation (*18*). Modeling the G1202R substitution based on the co-crystal structure of LA9 bound to wild-type ALK revealed that LA9 can maintain identical binding without significant steric clash, consistent with the nearly identical cellular IC_50_ values of LA9 against nonmutant and G1202R ALK. In contrast to G1202 which is positioned in the ALK solvent front, I1171 is buried within the ALK hydrophobic core, and I1171N/S/T mutations confer resistance by increasing catalytic activity of the kinase rather than steric hindrance of ligand binding (*23, 24*). Analysis of the co-crystal structure of LA7 bound to wild-type ALK revealed several interactions between the hydroxyl groups of the ligand and the D1203 side chain carboxylate and amino nitrogen, the G1201 backbone carbonyl and a structural water – all together forming a network of optimal interactions that strengthen ligand binding. Notably, the features of LA9 and LA7 that lead to improved selectivity against G1202R vs I1171N/S/T appear to be mutually exclusive. The increased functionality of the thiazole ring of LA7 that interacts with G1201 and D1203 becomes a liability in the presence of the G1202R mutation due to increased steric clash. Conversely, the lack of functionality on the phenyl ring of LA9 allows accommodation of the sterically bulky G1202R substitution, but prevents LA9 from forming similar favorable interactions with adjacent G1201 and D1203. Collectively, these findings provide a structure-based conceptual framework for different approaches to target distinct classes of *ALK* compound mutations that emerge following treatment with sequential second- and third-generation ALK inhibitors.

Given the current dearth of therapeutic options for patients after lorlatinib, several efforts to target lorlatinib-resistant compound *ALK* mutations are underway. Most notably, fourth-generation ALK inhibitors to target G1202R compound resistance mutations are being developed. As an example, TPX-0131 has demonstrated excellent potency against G1202R compound mutations in preclinical models (*26*). Reminiscent of our results, TPX-0131 appears to achieve improved activity through increased potency against the G1202R single mutation and does not target I1171 compound mutations, although the structural basis for selectivity has not been reported. In a similar fashion, NVL-655 harbors preclinical potency against G1202R compound mutations (*27*). Other studies have reported repurposing of available kinase inhibitors to overcome specific compound mutations. Okada *et al.* found that AG-957, initially developed as a BCR-ABL inhibitor, suppressed the viability of cells harboring G1202R+L1196M (IC_50_: 87nM) (*20*), while Mizuta et al. found that gilteritinib, a FLT3 inhibitor approved for the treatment of *FLT3*-mutant AML, could overcome I1171N compound mutations but not G1202R compound mutations (*28*). These reports together with our findings support that distinct compounds may be required in clinic to overcome G1202R- vs I1171N/S/T-based (vs other) *ALK* compound mutations.

Importantly, these results further underscore the challenges inherent to sequential targeted therapies and subsequent evolution of heterogeneous compound mutations. While our findings point to the G1202R- and I1171N/S/T-containing *ALK* compound mutations as the predominant lorlatinib-resistant mutations and therefore, the most urgent and salient target for drug development, we anticipate that not all of the broad array of *ALK* compound mutations that emerge in patients after lorlatinib will ultimately be surmountable, even with fourth-generation ALK inhibitors in development. Therefore, a more effective strategy may be to suppress the emergence of *ALK* mutations upfront. Our group previously showed in accelerated mutagenesis screening of cells expressing nonmutant EML4-ALK (modeling treatment-naïve tumors) that no single *ALK* mutation could confer resistance to lorlatinib, whereas in cells harboring pre-existing single *ALK* mutations (modeling tumors resistant to prior ALK TKIs), compound *ALK* mutations could emerge to confer lorlatinib resistance (*18*). Thus, upfront lorlatinib therapy may suppress, or at least delay, the development of *ALK* mutations. Recently, the interim analysis of the phase III CROWN study comparing front-line lorlatinib to crizotinib was published, demonstrating an impressive prolongation in the progression-free survival with the hazard ratio for disease progression or death of 0.28 (95% CI, 0.19-0.41) that was numerically the lowest among all phase III trials that evaluated next-generation ALK TKIs in the first-line setting (*7–9, 29*). On the basis of these results, the FDA approval of lorlatinib was expanded to include the first-line indication. While numerous considerations will ultimately factor into which next-generation ALK inhibitor is selected for first-line use in patients, one potential benefit of upfront lorlatinib may be in delaying or suppressing on-target resistance and circumventing the emergence of refractory compound *ALK* mutations, at least some of which may not be able to be overcome in patients. On the other hand, such an approach may result in the selection for ALK-independent resistance requiring combination strategies.

In summary, we have provided here an expanded catalogue of compound *ALK* mutations observed in patients treated with lorlatinib, highlighting G1202R- and I1171N/S/T-containing mutations as occurring most commonly after sequential therapy. We have shown through functional and structural analysis of a diverse set of lorlatinib analogs that distinct compounds may be required to target selective subsets of mutant ALK kinases, including G1202R- vs I1171N/S/T-harboring compound mutants. Our data offer a framework for the functional classification of heterogeneous compound mutations to guide the development of novel therapeutic strategies addressing on-target resistance after sequential targeted therapy.

## METHODS

### Patients and Tumor Genotyping

Patients were included if they had the diagnosis of advanced ALK-positive NSCLC, were treated at Massachusetts General Hospital (MGH), had disease progression on lorlatinib and underwent resistant tissue biopsies with genotyping after providing informed consent between November 2014 and October 2020. All cases underwent histopathology review on a clinical basis. Genotyping was performed using: the MGH SNaPshot DNA-based genotyping panel and a separate RNA-based NGS assay (Solid Fusion Assay) (n=35) as previously described (*30*), FoundationOne (n=9; Foundation Medicine, Inc.; Cambridge, MA), Caris NGS (n=1; Caris Life Sciences; Irving, TX), Tempus xT panel (n=1; Tempus; Chicago, IL), Oncomine Comprehensive Assay (n=1; Corneill, New York, NY), and MD Anderson Solid Tumor Genomic Assay (n=1; MD Anderson Cancer Center; Houston, TX).

Electronic medical records were retrospectively reviewed to extract data on clinical, pathologic, and molecular features. Data were updated as of January 2021. Time to progression (TTP) was measured from the time of therapy initiation to clinical/radiologic disease progression. Time to treatment discontinuation (TTD) was measured from the time of lorlatinib initiation to lorlatinib discontinuation. Patients continuing on treatment at the time of data cut-off were censored at their last follow-up. The study was performed under an Institutional Review Board–approved protocol. This study was conducted in accordance with the Belmont Report and the U.S. Common Rule.

### Plasma Biopsy Dataset

We queried a de-identified dataset at Guardant Health (Redwood City, CA) comprised of plasma specimens analyzed between October 2015 and December 2020 in order to identify specimens harboring ≥2 *ALK* kinase domain mutations. All plasma specimens were genotyped using the Guardant360 cell-free DNA (cfDNA) assay, as previously described (*31*). The Guardant360 assay detects *ALK* fusions in addition to mutations in critical exons of ALK, including exons 22-25 which encode the ALK kinase domain. The allelic configuration of *ALK* mutations (i.e., *cis* vs *trans*) was determined for a subset of specimens with multiple *ALK* mutations arising in the same exon.

### Cell Lines and Reagents

Ba/F3 immortalized murine bone marrow-derived pro-B cells were obtained from the RIKEN BRC Cell Bank (RIKEN BioResource Center) in 2010 and cultured in DMEM with 10% FBS with (parental) or without (EML4-ALK) IL3 (0.5 ng/mL). cDNAs encoding EML4-ALK variant 1 (E13; A20) and variant 3a (E6a; A20) containing different point mutations were cloned into retroviral expression vectors, and Ba/F3 cells were infected with the virus as described previously (24). After retroviral infection, Ba/F3 cells were selected in puromycin (0.7 μg/mL) for 2 weeks. IL3 was withdrawn from the culture medium for more than 2 weeks before experiments.

The MGH953-4 cell line was developed from a malignant pleural effusion from a patient with ALK-positive NSCLC who had progressed on alectinib, and the MGH953-7 cell line was subsequently developed from a patient-derived xenograft (PDX) model established from a lorlatinib-resistant malignant pleural effusion from the same patient. The MGH990-2 cell line was developed from an adrenal biopsy from a patient with lorlatinib-resistant ALK-positive NSCLC. Prior to cell line generation, the patients signed informed consent to participate in a Dana Farber/Harvard Cancer Center Institutional Review Board-approved protocol giving permission for research to be performed on their sample. MGH953-4, MGH953-7, and MGH990-2 were cultured in RPMI1640 with 10% FBS. Cell lines were sequenced to confirm the presence of ALK rearrangement and ALK mutations. Additional authentication was performed by SNP fingerprinting. The expression vectors of EML4-ALK were transfected with TransIT®-LT1 Transfection Reagent (Mirus Bio, LLC) according to the manufacturer’s protocol. Crizotinib and ceritinib were purchased from Selleck Chemicals. Alectinib and brigatinib were purchased from MedChem Express. Lorlatinib and lorlatinib analogs were provided by Pfizer, Inc.

### Mutagenesis

QuickChange XL Site-Directed mutagenesis kit (Agilent Technologies) was used to generate point mutations in the expression vectors of EML4-ALK in accordance with the manufacturer’s protocol.

### Survival Assays

Ba/F3 cells (2,000) or patient derived cells (5,000) were plated in triplicate into 96-well plates. Cells were incubated with CellTiter-Glo (Promega) after 48 or 72 hours after drug treatment, and luminescence was measured with a SpectraMax M5 Multi-Mode Microplate Reader (Molecular Devices, LLC). GraphPad Prism (GraphPad Software) was used to graphically display data and determine IC_50_ values by a nonlinear regression model utilizing a four-parameter analytic method.

### Western Blot Analysis

Ba/F3 cells and patient derived cell lines were treated for 6 hours with lorlatinib, LA7 or LA9 unless otherwise noted. Total protein lysates were analyzed by western blotting with the following antibodies (all from Cell Signaling Technology): phospho-ALK Y1604 (3341), phospho-ALK Y1282/83 (9687), ALK (3633), phospho-AKT S473 (4060), AKT (4691), phospho-ERK1/2 T202/Y204 (9101), ERK1/2 (9102), phospho-S6 S240/244 (5364), S6 (2217), and β-Actin (4970).

### Development of Xenografts and *in vivo* Pharmacological Studies

All animal studies were conducted in accordance with the guidelines as published in the Guide for the Care and Use of Laboratory Animals and were approved by the Institutional Animal Care and Use Committee of MGH. Female NSG mice aged 6 to 8 weeks were obtained from Jackson Laboratory. Mice were maintained in laminar flow units in sterile filter-top cages with Alpha-Dri bedding. MGH953-7 cells were implanted subcutaneously into the flank of mice. Similarly, 5 × 10^6^ 3T3-ALK I1171N+D1203N cells in 0.2 mL 50% BD Matrigel Basement Membrane Matrix in PBS were subcutaneously injected into the flank of mice. Mice were randomized into groups once the tumors had attained a volume of 200 mm^3^. The treatment groups were treated 5 days per week with drug solution dissolved in acid water (lorlatinib and LA9) or water (LA7) by oral gavage. Tumor volumes were measured twice weekly and calculated using the formula: mm^3^ = 0.52 x L x W^2^. For pharmacodynamics studies, tumor-bearing mice were administered drugs daily for 3 days, and tumors were harvested 3 hours after the last treatment for western blot and immunohistochemistry.

### Immunohistochemistry

Tissue sections 4 μm thick were cut from 10% buffered formalin-fixed and paraffin-embedded (FFPE) mice tumor tissue blocks. The deparaffinized and rehydrated sections were autoclaved in Bond TM Epitope Retrieval 2-1L EDTA (pH 9.0) at 100°C for 20 minutes for antigen retrieval. They were then incubated with rabbit polyclonal anti-phospho ALK (Y1507) antibody diluted 1:50 (ABCAM™) for 15 minutes at room temperature. Immunoactivity was detected with mixed DAB refine after 10 minutes (Leica Biosystems). The sections were counterstained with hematoxylin for 5 minutes and dehydrated prior to coverslipping.

### Pharmacokinetic Analysis

During the *in vivo* pharmacology studies, serial whole blood samples (~8 uL/time point) were collected from the tail vein of mice (n = 3/group) at 0 (pre-dose),1, 4, 7, and 24 hours after the last dose of a 5-day or longer treatment period. The concentration of lorlatinib, LA7 or LA9 in whole blood was determined using quantitative LC-MS/MS methods. Pharmacokinetic parameters for each analyte were calculated from individual animal concentration-time profiles using standard non-compartmental methods. The unbound fraction (f_u,p_) of each analyte in NOD-scid mouse plasma was determined *in vitro* using an equilibrium dialysis method as described previously (*32*). The blood-to-plasma concentration ratio (BPR) of each analyte was determined *in vitro* in fresh whole mouse (CD-1) blood as described previously (*32*). The unbound fraction in blood (f_u,b_) for each analyte was calculated from their respective measured f_u,p_ and BPR values as follows: f_u,b_ = f_u,p_ / BPR.

### Structural Modeling of Lorlatinib Analogs Bound to mutant ALK

Co-crystal structures of wild-type ALK were used as the starting point for modelling mutations. The crystal structural data and methods have been deposited to the Protein Data Bank (PDB) and have been assigned ID codes: lorlatinib, PDB 4CLI (*14*); LA7, PDB 7R7R; LA9, PDB 7R7K. Mutations were introduced and side chains minimized using the OPLS2003 forcefield. Protein and ligand minimizations were also performed with flexible protein using OPLS2003 forcefield. Original and protein-ligand minimized structures were overlapped to visualize extent of protein movement, which inversely correlates with fit from original.

### Statistical Analysis

TTP and TTD curves were estimated using the Kaplan-Meier method with the differences between groups expressed as a hazard ratio (HR) and compared using the log-rank test. The Cox proportional hazards model was used to estimate the HR for TTP, while the HR for TTD was estimated as a ratio of the observed group medians under the assumption of an exponential distribution as none of the patients with known baseline *ALK* mutations had discontinued treatment earlier than any of those without known baseline *ALK* mutations. All statistical analyses were performed using Stata version 14.2.

## ACKNOWLEDGEMENTS

We thank patients and their families, as well as members of the Hata lab and MGH Thoracic Oncology Group for helpful discussion and support. This study was supported by a JSPS Overseas Research Fellowships (to AS), a National Cancer Institute Career Development Award (K12CA087723–16 to IDJ), NIH/NCI R01CA164273 (to JJL), a Lung Cancer Research Foundation grant (to SY), by Be a Piece of the Solution, and by Targeting a Cure for Lung Cancer Research Fund at MGH.

## COMPETING INTERESTS STATEMENT

**TWJ, TRJ, PW, SLW** and **MAM** are employees of Pfizer, Inc. **IDJ** has received honoraria from Foundation Medicine, Creative Education Concepts, and American Lung Association, consulting fees from Guidepost, AstraZeneca, Boehringer Ingelheim, BostonGene, Catalyst, Novocure, Pfizer, Syros, and Xcovery, research support from Array, Genentech, Novartis, Pfizer, and Guardant Health, and travel support from Array and Pfizer. **LAK** is an employee and shareholder of Guardant Health. **JFG** has served as a compensated consultant or received honoraria from Bristol-Myers Squibb, Genentech, Ariad/Takeda, Loxo, Pfizer, Incyte, Novartis, Merck, Agios, Amgen, Jounce, Karyopharm, GlydeBio, Regeneron, Oncorus, Helsinn, Jounce, Array, and Clovis Oncology, has an immediate family member who is an employee with equity in Ironwood Pharmaceuticals, has received research funding from Novartis, Genentech/Roche, and Ariad/Takeda, and institutional research support from Tesaro, Moderna, Blueprint, BMS, Jounce, Array, Adaptimmune, Novartis, Genentech/Roche, Alexo and Merck. **JJL** has served as a compensated consultant for Genentech, C4 Therapeutics, Blueprint Medicines, Nuvalent, Turning Point Therapeutics, and Elevation Oncology; received honorarium and travel support from Pfizer; received institutional research funds from Hengrui Therapeutics, Turning Point Therapeutics, Neon Therapeutics, Relay Therapeutics, Bayer, Elevation Oncology, Roche/Genentech, Pfizer, and Novartis; received CME funding from OncLive, MedStar Health, and Northwell Health. **SY** has received a consulting fee from Pfizer Japan. **ANH** has received research support from Pfizer, Nuvalent Inc., Roche/Genentech, Amgen, Blueprint Medicines, Eli Lilly, Relay Therapeutics and Genentech; has served as a paid consultant for Nuvalent.

---

**Supplementary Table 1.** Baseline characteristics of patients with advanced ALK-positive lung cancer included in the tissue lorlatinib resistance cohort.

| **Patient characteristic** | **n (N=47)** | **%** |
| --- | --- | --- |
| **Age at diagnosis,\* median (range)** | 49 | (22-69) |
| **Gender** |  |  |
| Male | 18 | 38.3 |
| Female | 29 | 61.7 |
| **Race** |  |  |
| White | 38 | 80.9 |
| Asian | 5 | 10.6 |
| Other | 4 | 8.5 |
| **Smoking history** |  |  |
| Never or light smoker | 43 | 91.5 |
| Heavy smoker | 4 | 8.5 |
| **Histology** |  |  |
| Adenocarcinoma | 46 | 97.9 |
| Other | 1 | 2.1 |
| **ALK fusion** |  |  |
| EML4-ALK v1 | 11 | 23.4 |
| EML4-ALK v3 | 23 | 48.9 |
| EML4-ALK v2 | 3 | 6.4 |
| Other EML4-ALK | 5 | 10.6 |
| Non-EML4-ALK | 2 | 4.3 |
| Unknown | 3 | 6.4 |
| **Prior ALK TKI** |  |  |
| Prior crizotinib + 2nd-generation TKI(s) | 36 | 76.6 |
| Prior 2nd-generation TKI(s), no criz | 11 | 23.4 |
| **Number of prior distinct ALK TKI(s)** |  |  |
| One<sup>§</sup> | 7 | 14.9 |
| Two | 23 | 48.9 |
| Three | 15 | 31.9 |
| Four | 2 | 4.3 |

\*Age at diagnosis of advanced/metastatic disease
<sup>§</sup>All of these patients received one prior second-generation ALK inhibitor

**Supplementary Table 2.**
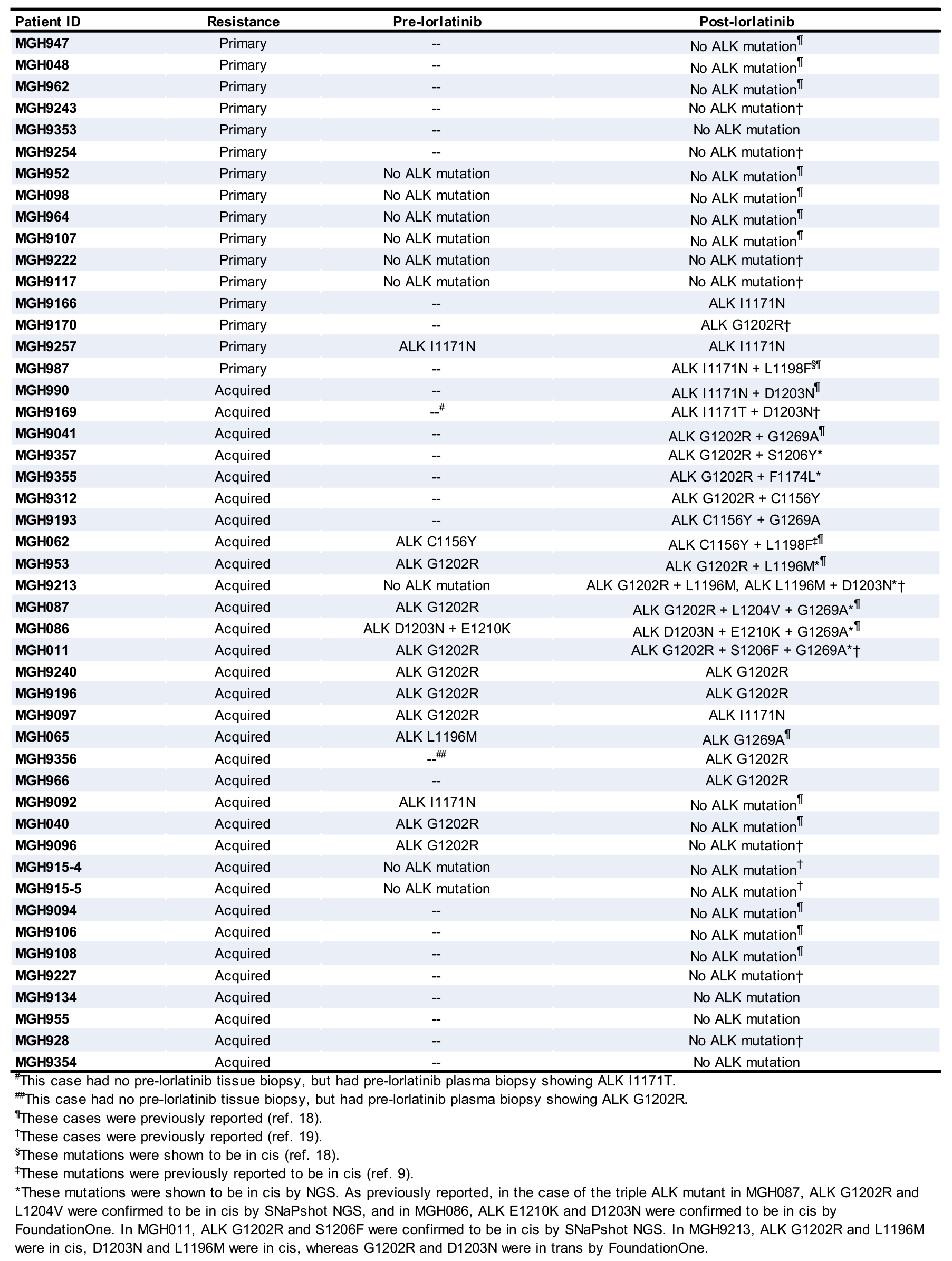
*ALK* resistance mutations detected in lorlatinib-resistant tissue biopsies. The baseline *ALK* mutation detected prior to lorlatinib treatment (‘pre-lorlatinib’) is shown where known (‘--’ indicates that pre-lorlatinib tissue analysis was not performed).

**Figure S1.**
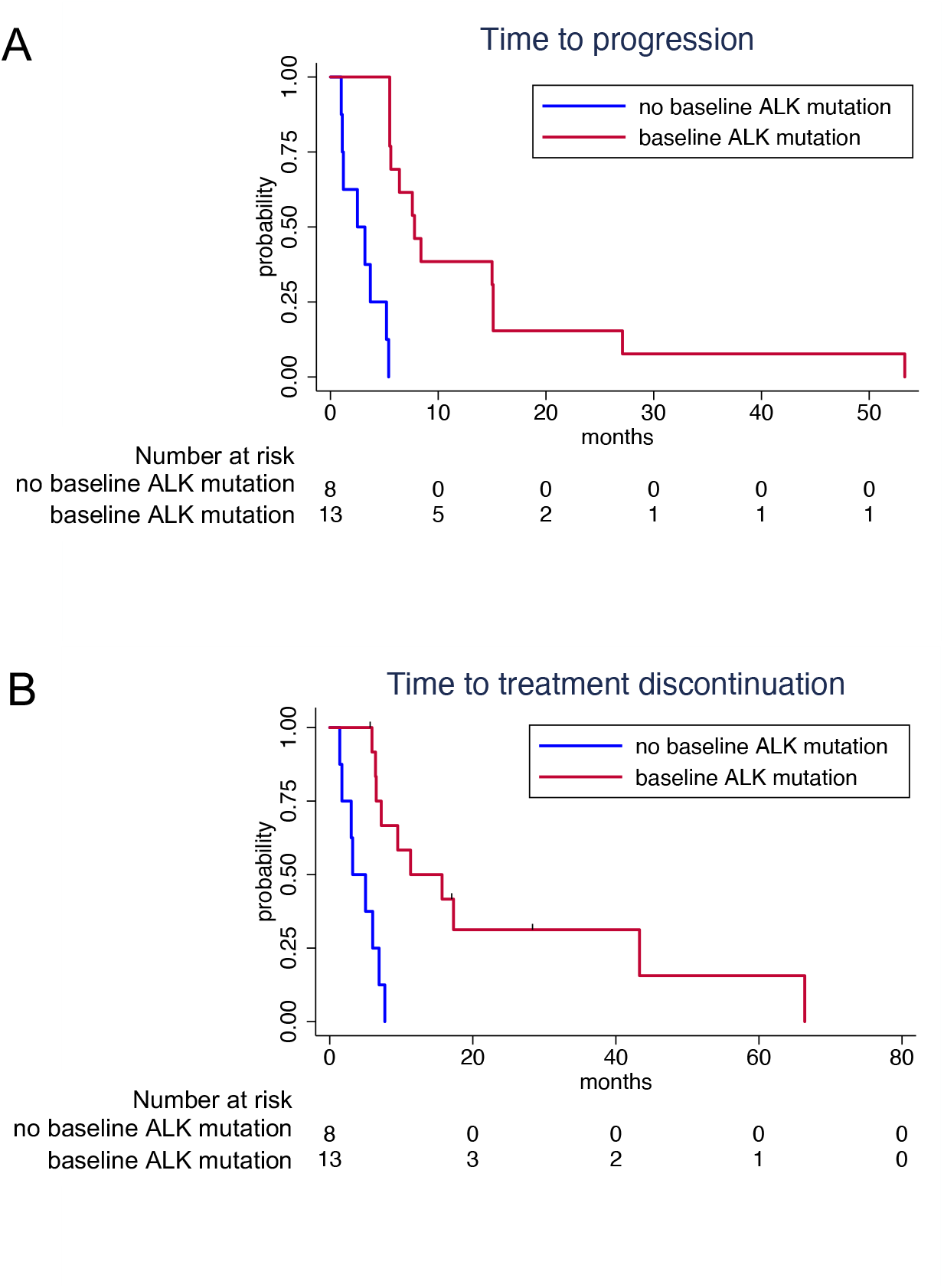
Time to progression on lorlatinib (A) and time to treatment discontinuation from lorlatinib (B) according to the presence or absence of a baseline *ALK* mutation. Only those patients with available pre-lorlatinib tissue biopsy and known status of baseline *ALK* mutations are included in these analyses.

**Figure S2.**
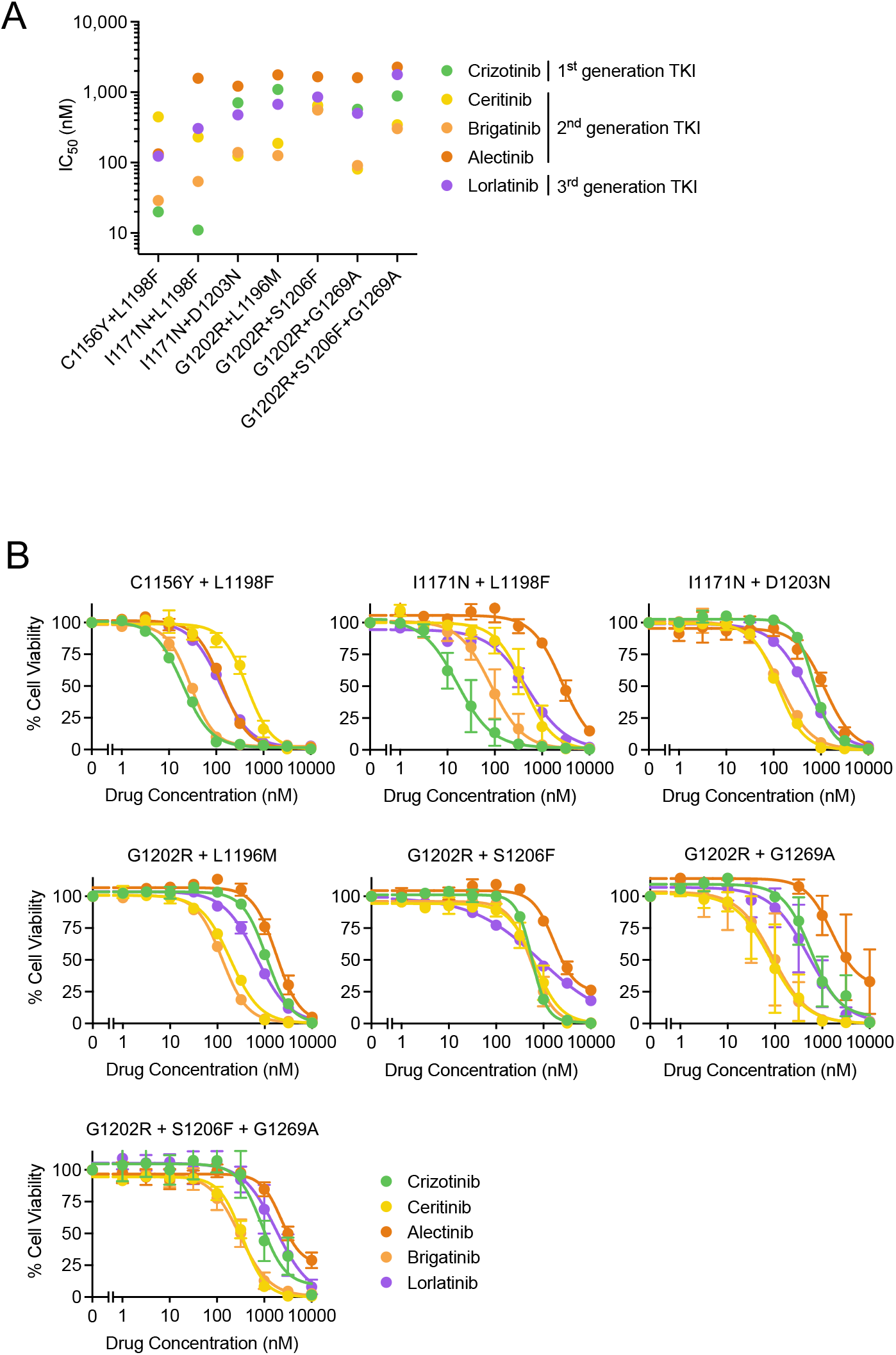
Sensitivity of compound *ALK* mutations to currently FDA-approved ALK tyrosine kinase inhibitors. Cellular IC_50_ values (A) and dose response curves (B) of Ba/F3 cells expressing clinical *ALK* compound mutantations to crizotinib, ceritinib, brigatinib, alectinib, and lorlatinib. Data correspond to the heatmap shown in Fig. 1E. Error bars indicate SE.

**Figure S3.**
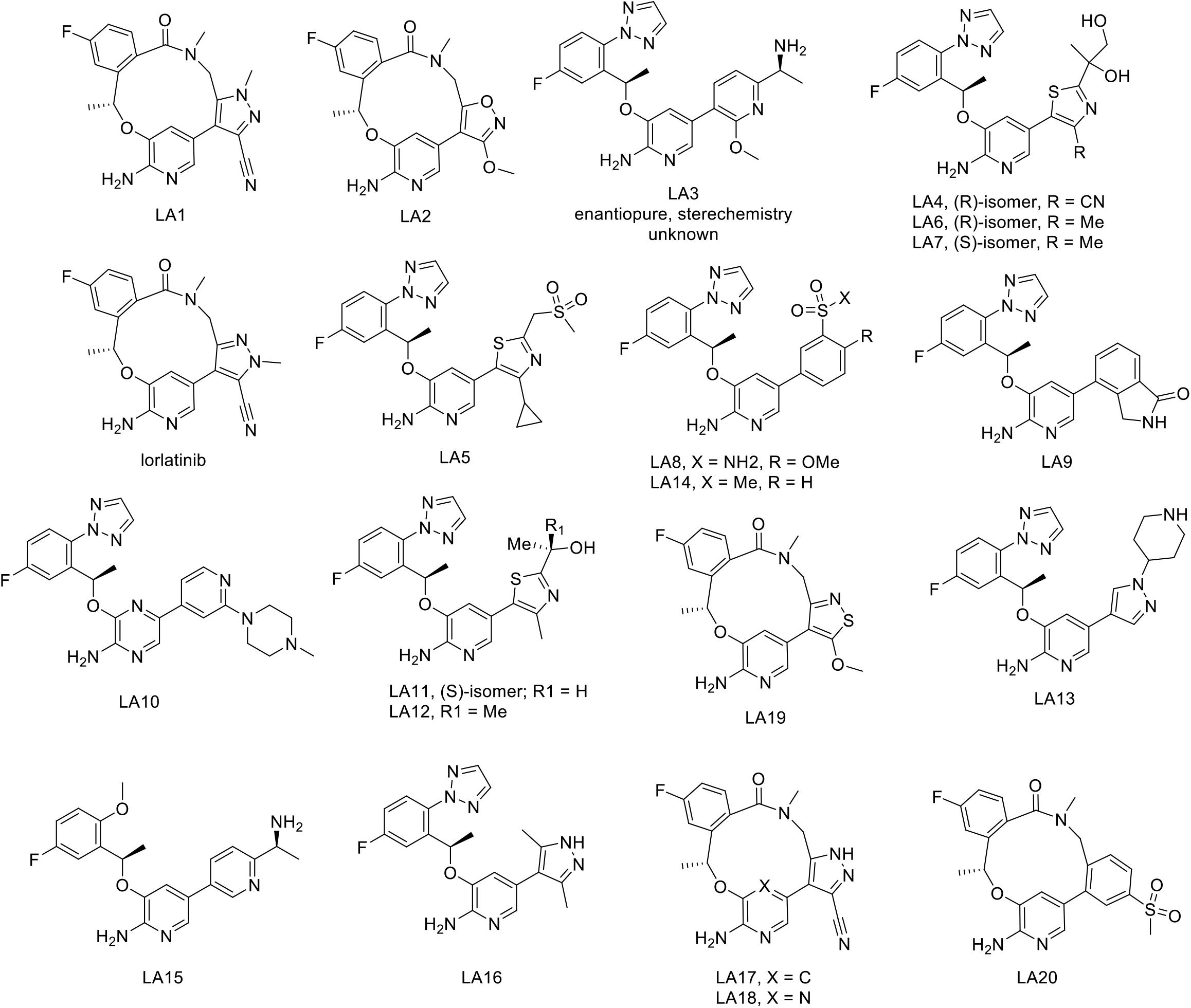
Molecular structures of lorlatinib analogs (LAs) 1-20.

**Figure S4.**
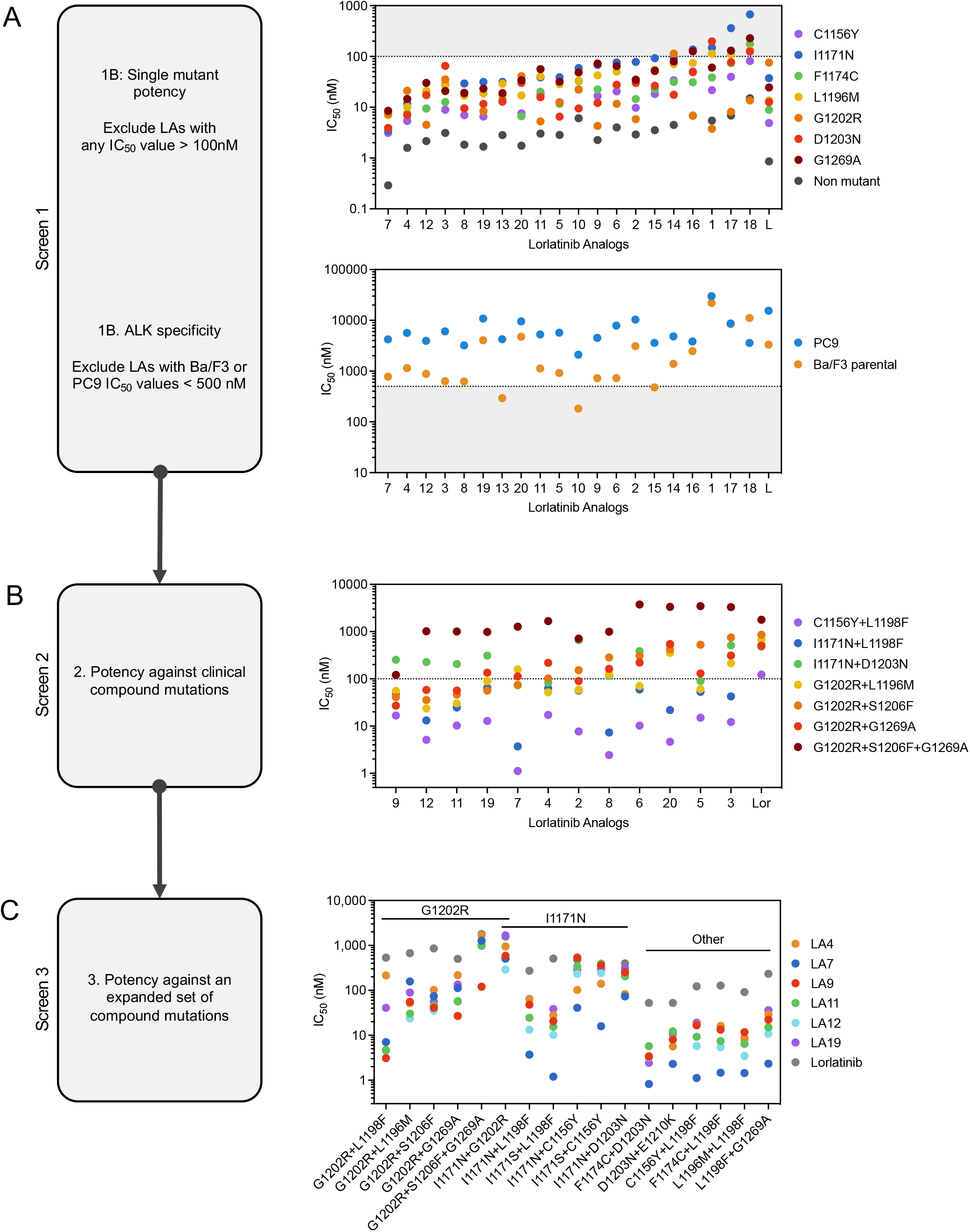
Drug screening with lorlatinib analogs using Ba/F3 *ALK* mutation models. (A) Cellular IC_50_ values of each LA against single *ALK* mutations (top) and Ba/F3 parental and PC9 cells (bottom). This data corresponds to the heatmap in Figure 1B. 12 LAs were selected for further validation. (B) Cellular IC_50_ values of 12 LAs against clinical *ALK* compound mutations. This data corresponds to the heatmap in Figure 1C. 6 LAs were selected for further validation. (C) Cellular IC_50_ values of 6 LAs against an expanded set of clinical and putative compound *ALK* mutations. This data corresponds to the heatmap in Figure 1D.

**Figure S5.**
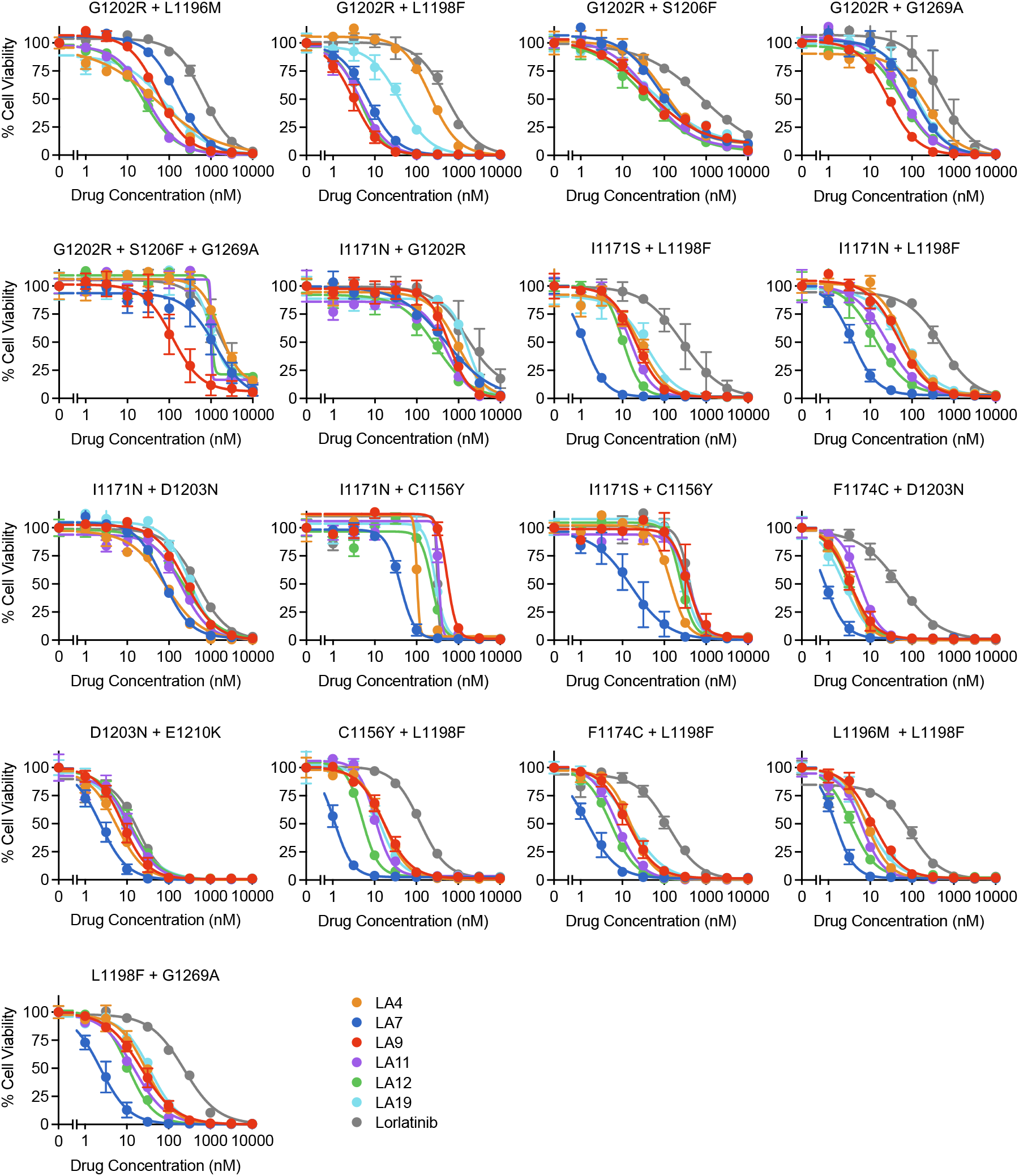
Drug dose response curves of lorlatinib analogs against Ba/F3 *ALK* mutation models. Cell viability assays performed with Ba/F3 cells expressing EML4-ALK with the indicated compound mutations treated with 6 LAs or lorlatinib for 48 hours. The viabilities were measured with CellTiter-Glo assay. These data correspond to the heatmap in Figure 1D and dot plots in Figure S4C. Error bars indicate SE.

**Figure S6.**
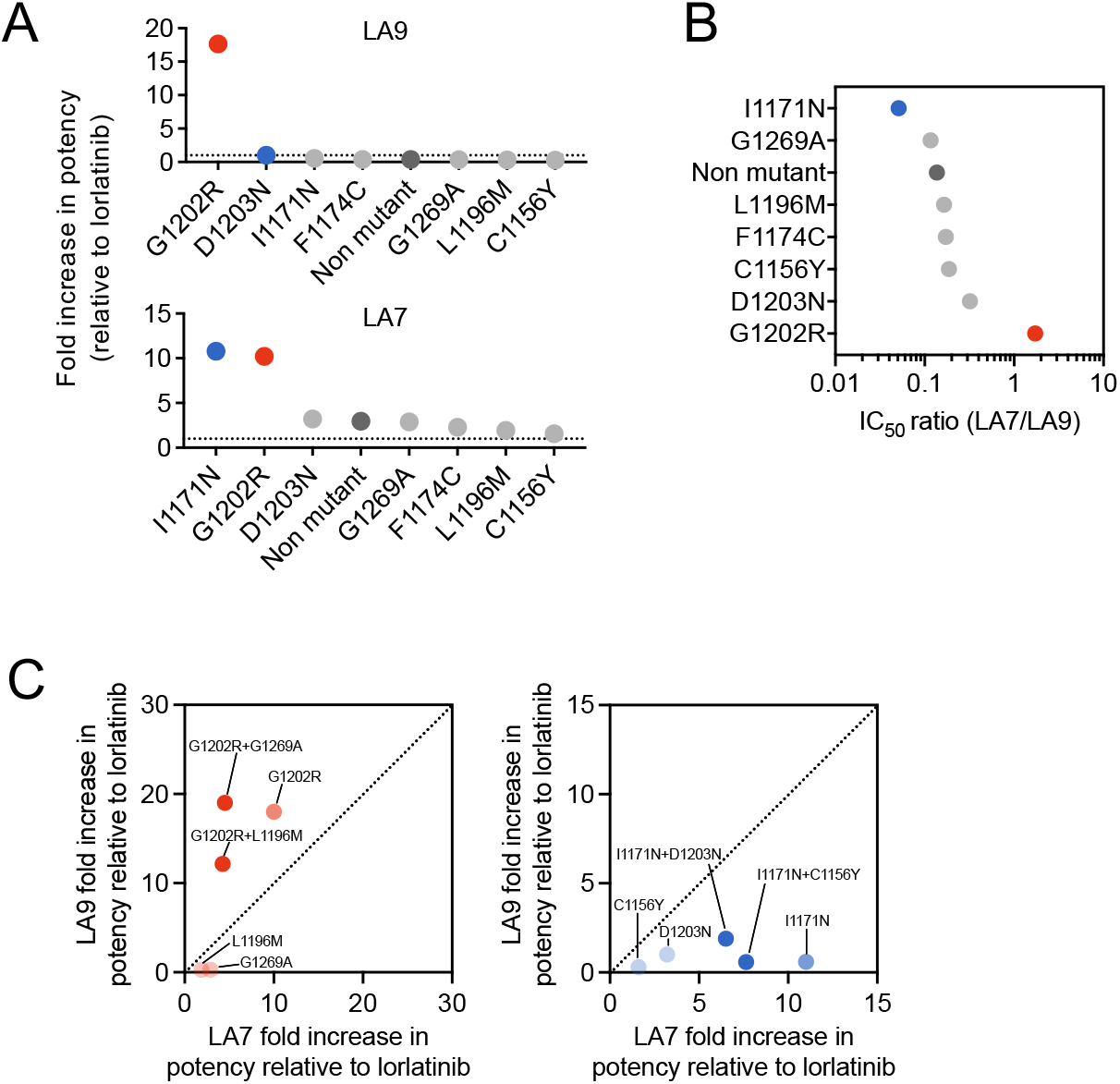
Selectivity of LA7 and LA9 against I1171N and G1202R single and compound mutants, respectively. (A) Increase in potency of LA9 and LA7 against single ALK mutants relative to lorlatinib, calculated by the ratio IC_50_(lorlatinib)/IC_50_(LA). (B) Relative selectivity of LA7 and LA9 for single ALK mutants, calculated by the ratio IC_50_(LA7)/IC_50_(LA9). (C) Comparison of relative fold increase in potency of LA7 vs LA9 (over lorlatinib) against single and compound ALK mutants. LA9 has a greater increase in potency relative to lorlatinib against G1202R single and compound mutations compared to LA7 (left panel). Conversely, LA7 has greater increase in potency relative to lorlatinib against I1171N single and compound mutations compared to LA9 (right panel).

**Figure S7.**
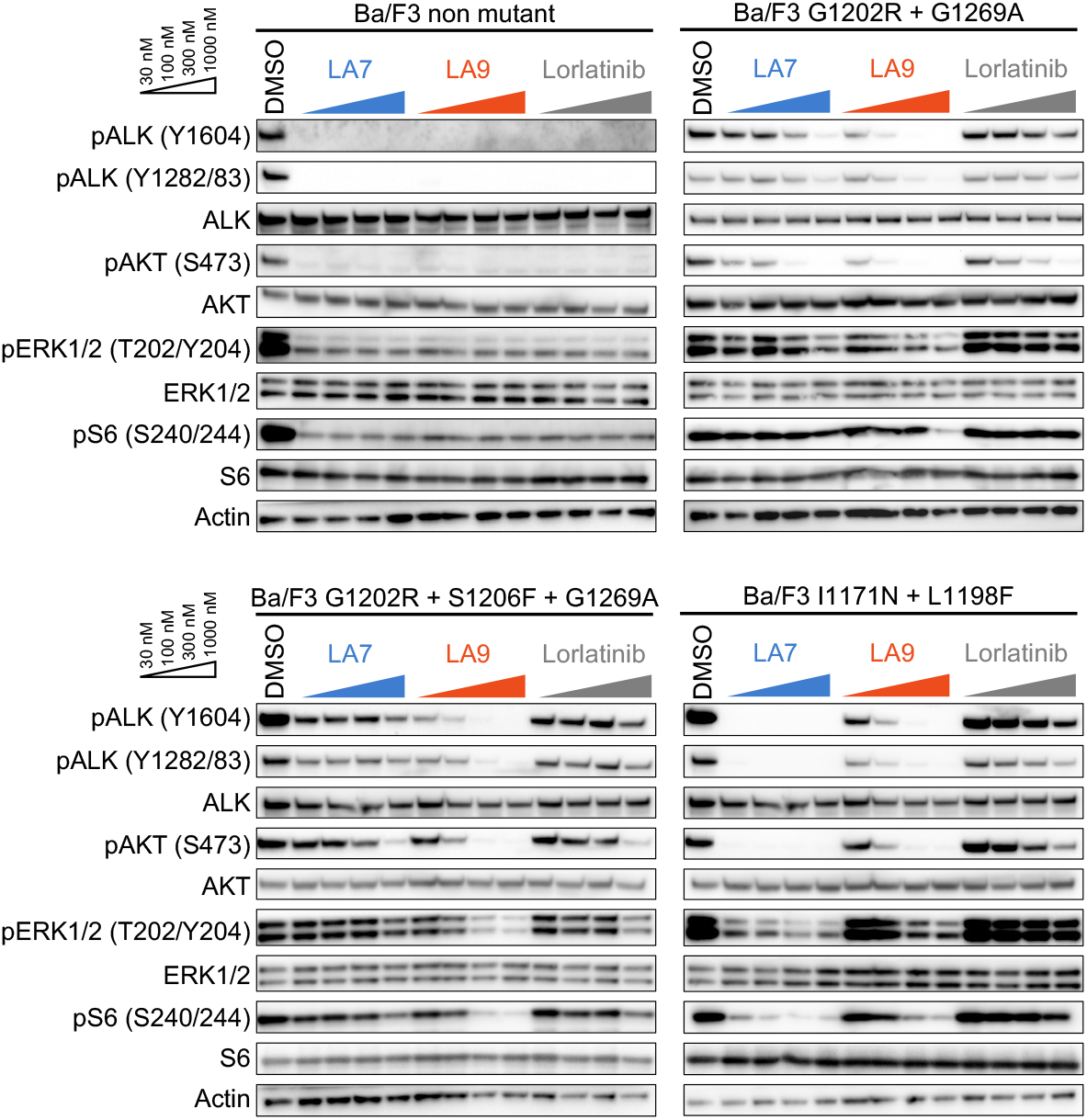
Western blot analysis performed on compound mutation Ba/F3 models. Ba/F3 cells expressing nonmutant EML4-ALK, EML4-ALK G1202R+G1269A, G1202R+S1206F+G1269A or I1171N+L1198F were treated with DMSO, LA7, LA9, or lorlatinib for 6 hours, and total cell lysates were analyzed by western blotting.

**Figure S8.**
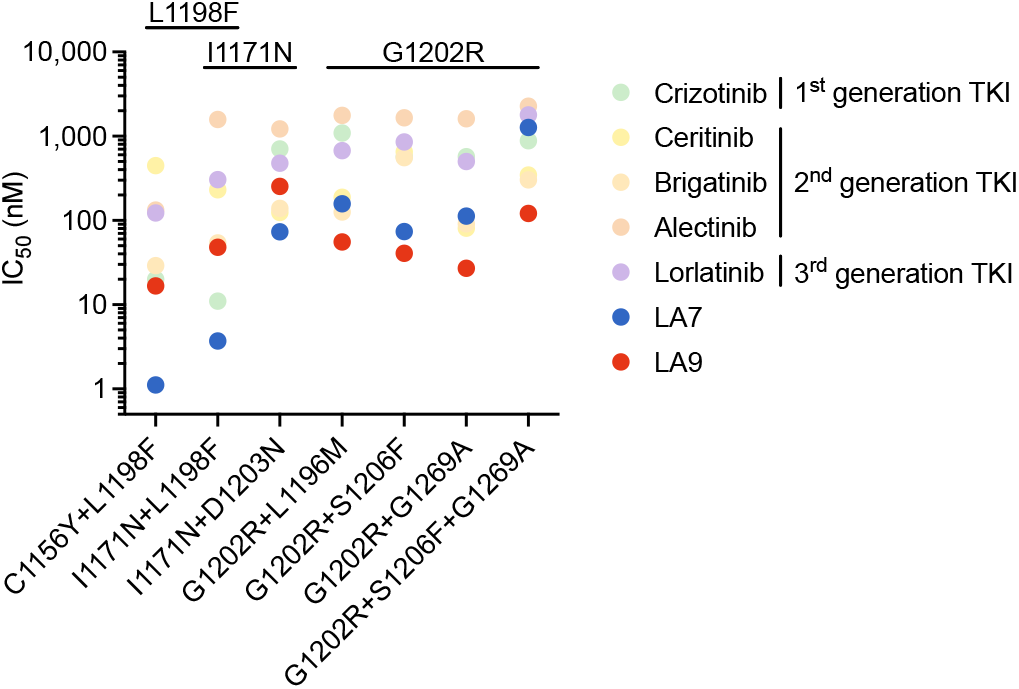
Comparison of LA7 and LA9 with currently approved ALK inhibitors. Cellular IC_50_ values of LA7, LA9 and currently approved ALK inhibitors including lorlatinib against clinical ALK compound mutations. IC_50_ values for approved ALK inhibitors are replotted from Figure S2 for comparison purposes.

**Figure S9.**
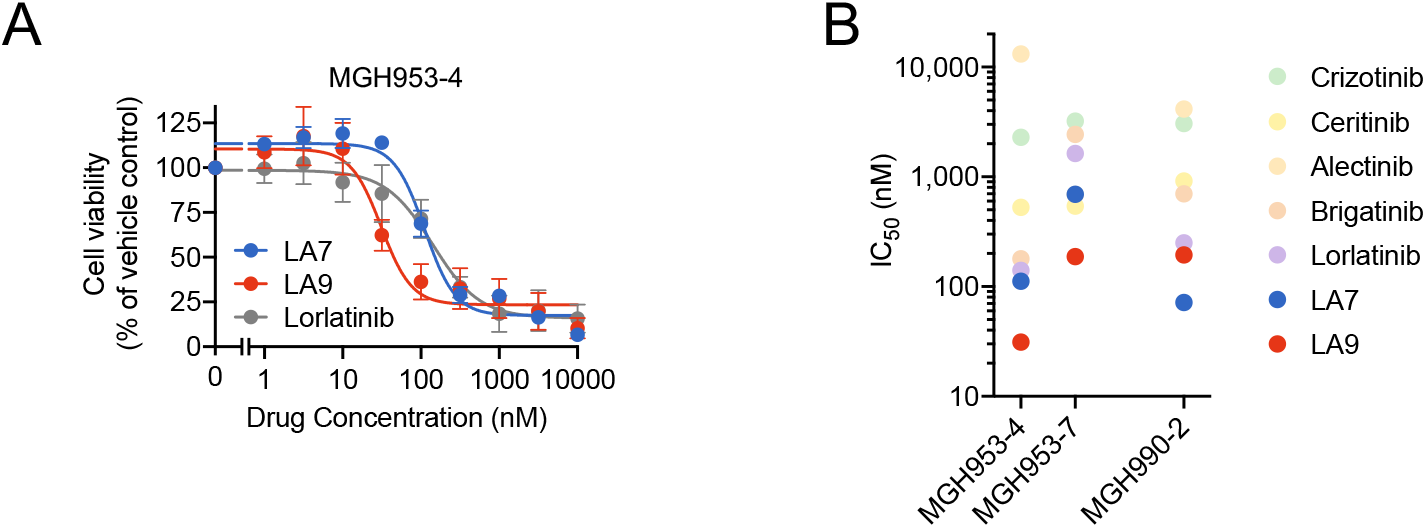
Activity of LA7 and LA9 in patient-derived models. (A) MGH953-4 (G1202R) cells were treated with the indicated drugs for 72 hours and cell viability assessed with CellTiter-Glo assay. Error bars indicate SE. (B) Cellular IC_50_ values of LA7 and LA9 compared to approved ALK inhibitors in MGH953-4, MGH953-7 (G1202R+L1196M), and MGH990-2 (I1171N+D1203N) cells.

**Figure S10.**
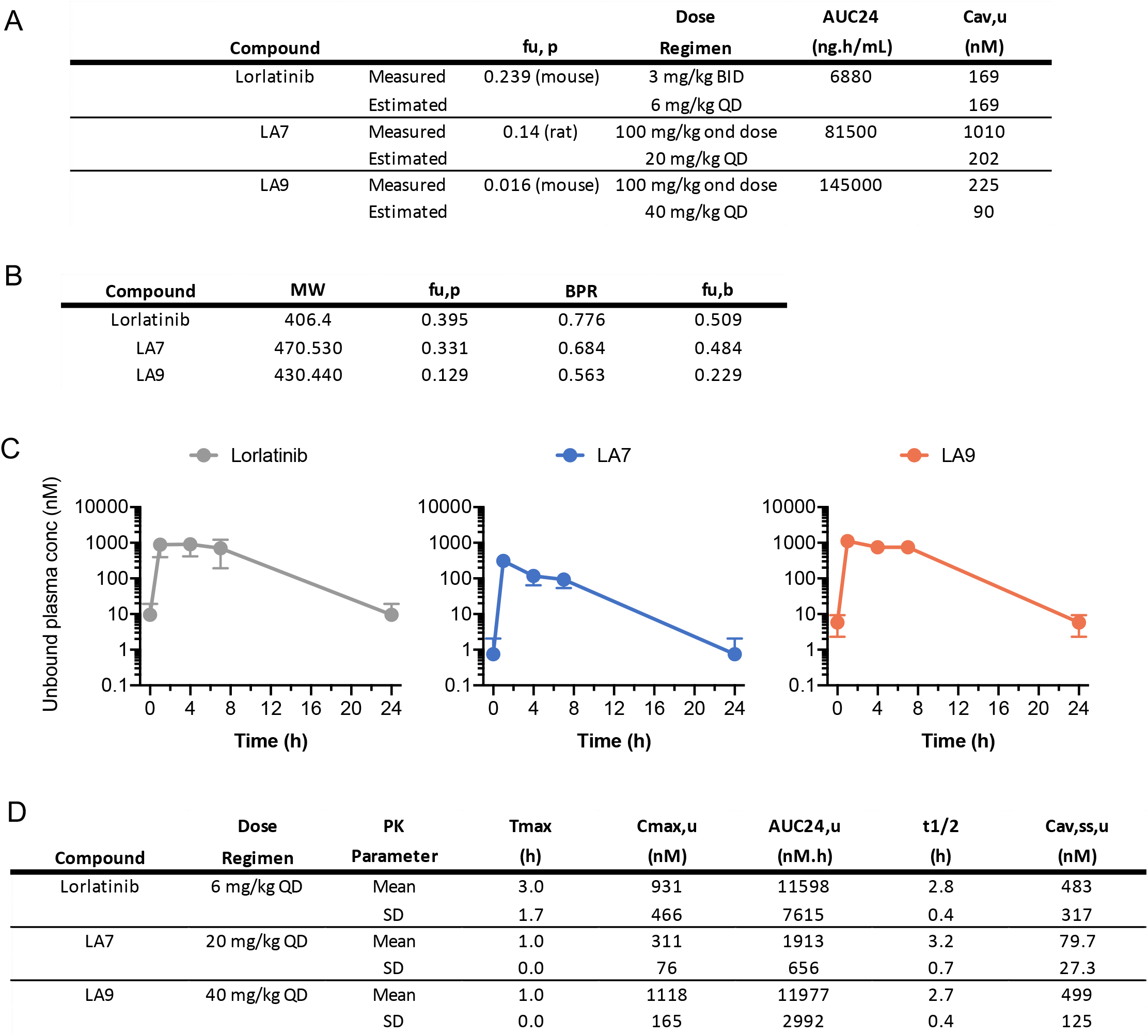
Pharmacokinetic parameters of lorlatinib, LA7, LA9. (A) To estimate the unbound plasma concentrations of lorlatinib, LA7 and LA9 to select comparable doses using a once-daily (QD) dosing regimen, we used available (historical) unbound fraction data of these drugs in either mouse or rat plasma and the AUC24 of lorlatinib at 3 mg/kg BID, LA7 at 100 mg/kg QD or LA9 at 100 mg/kg QD in mice. We estimated the average unbound plasma concentration for each drug to achieve approximately similar unbound systemic exposure: (1) lorlatinib: 169 nM at 6 mg/kg QD; (2) LA7: 202 nM at 20 mg/kg QD; and (3) LA9: 90 nM at 40 mg/kg QD. (B) During the *in vivo* pharmacology studies, serial, whole blood samples were collected at 0, 1, 4, 7 and 24 hours after the last dose for the determination of steady-state systemic exposure of each drug (*n* = 3 mice per group). To enable comparison of unbound systemic exposure of each drug *in vivo* in mice, plasma protein binding (f_u,p_) and blood-to-plasma concentration ratio (BPR) values were measured *in vitro* in mouse (NOD-scid) plasma and mouse whole blood, respectively. Plasma protein binding values determined in NOD-scid mice for lorlatinib and LA7 were generally consistent with the initially available (historical) values in mouse (CD-1) or rat plasma, whereas LA9 was approximately 10-fold less bound relative to the initially available f_u,p_ value determined previously in CD-1 mouse plasma. The apparent discrepancy between these f_u,p_ values could be due to a strain difference in the extent of plasma protein binding or differences and/or experimental variability in the respective *in vitro* protein binding assays utilized. (C) Unbound systemic concentrations of lorlatinib, LA7 and LA9 were calculated based on the total drug concentrations in whole blood measured *in vivo* and the unbound fraction in blood (f_u,b_; calculated as f_u,p_ / BPR). Error bars indicate SE. (D) The mean unbound average plasma concentrations at steady-state (C_av,ss,u_) of lorlatinib and LA9 were similar (483 nM or 499 nM, respectively), with a lower value observed for LA7 (79.7 nM). Abbreviations: f_u,p_ (unbound fraction in plasma), AUC_24_ (area under the total plasma concentration-time curve from 0 to 24 hr), C_av,ss,u_ (average unbound plasma concentration at steady-state, or AUC_24_ / 24 hr), MW (molecular weight), BPR (blood-to-plasma concentration ratio), f_u,b_ (unbound fraction in blood, or f_u,p_ / BPR), T_max_ (time to reach maximum plasma concentration following drug administration), C_max,u_ (maximum unbound plasma concentration), AUC_24,u_ (unbound plasma AUC_24_, or AUC_24_ × f_u,p_).

**Figure S11.**
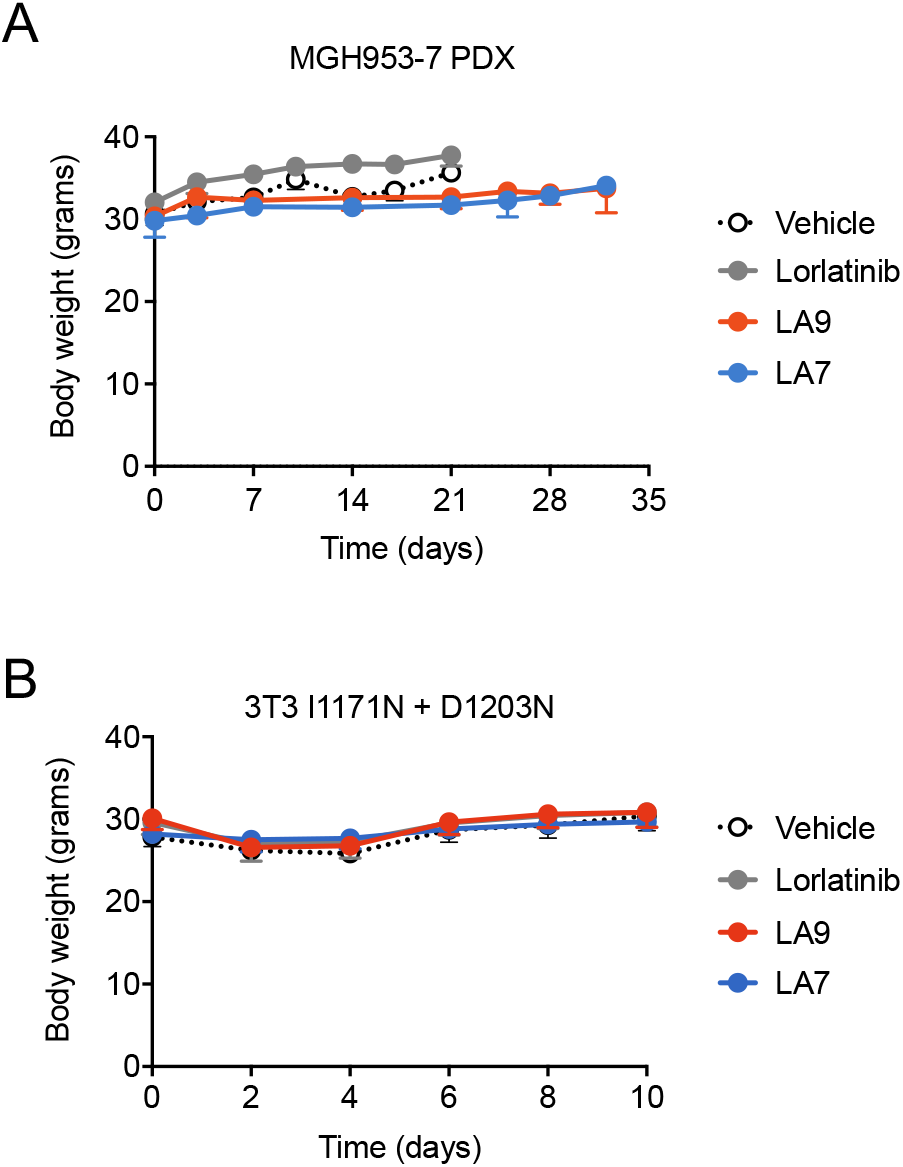
Body weight of mice treated with lorlatinib and LAs. Body weight of mice bearing MGH953-7 (A) and NIH3T3 EML4-ALK I1171N+D1203N (B) xenograft tumors during the course of treatment. Error bars indicate SE.

**Figure S12.**
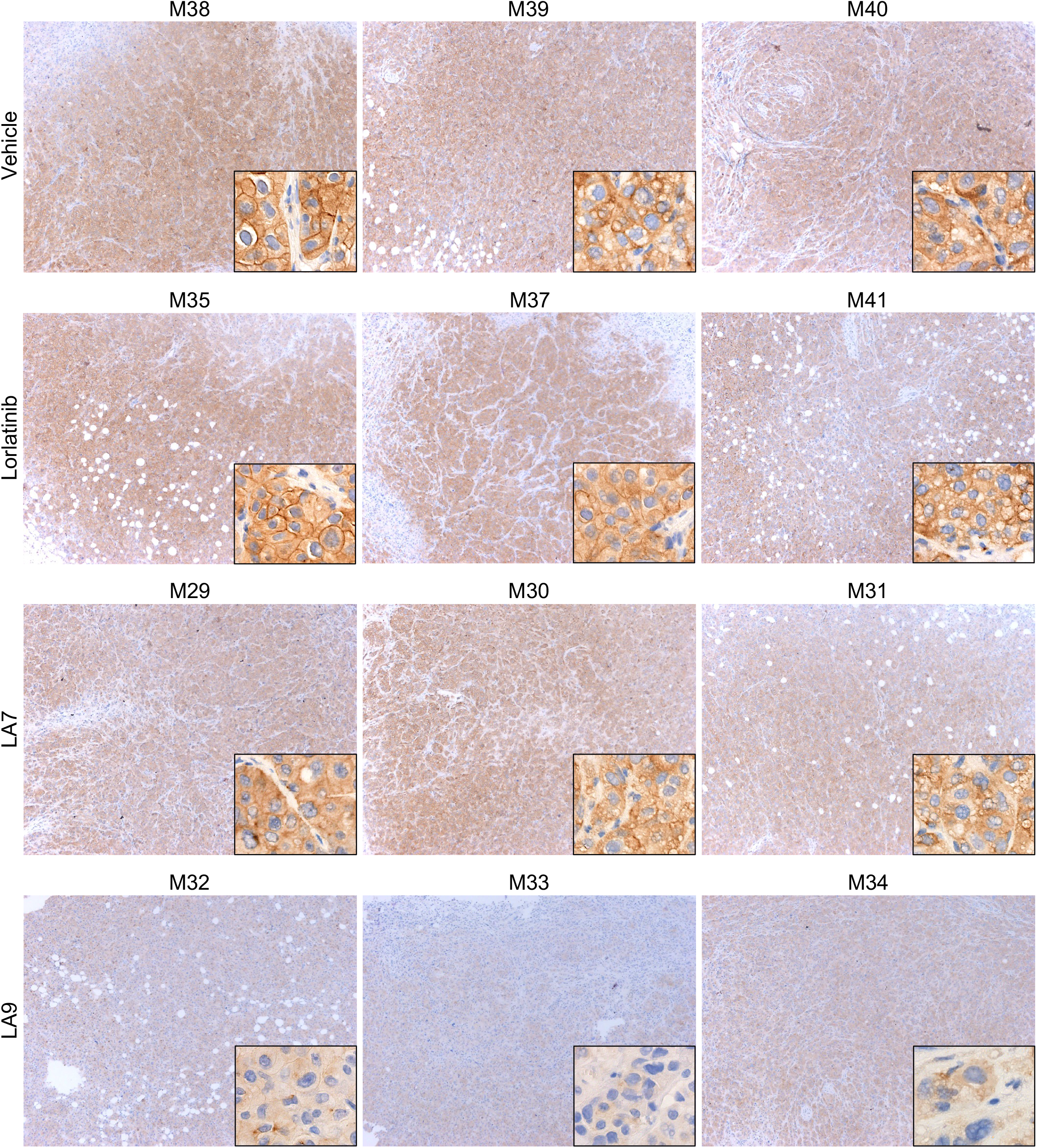
Phospho-ALK immunohistochemistry of MGH953-7 PDX tumors. MGH953-7 PDX tumors treated with lorlatinib, LA7 or LA9 were harvested after 3-day treatment and FFPE sections were stained with anti-phospho-ALK antibody. Phospho-ALK is localized in the cytoplasm and exhibits a diffuse staining pattern in tumor cells.

**Figure S13.**
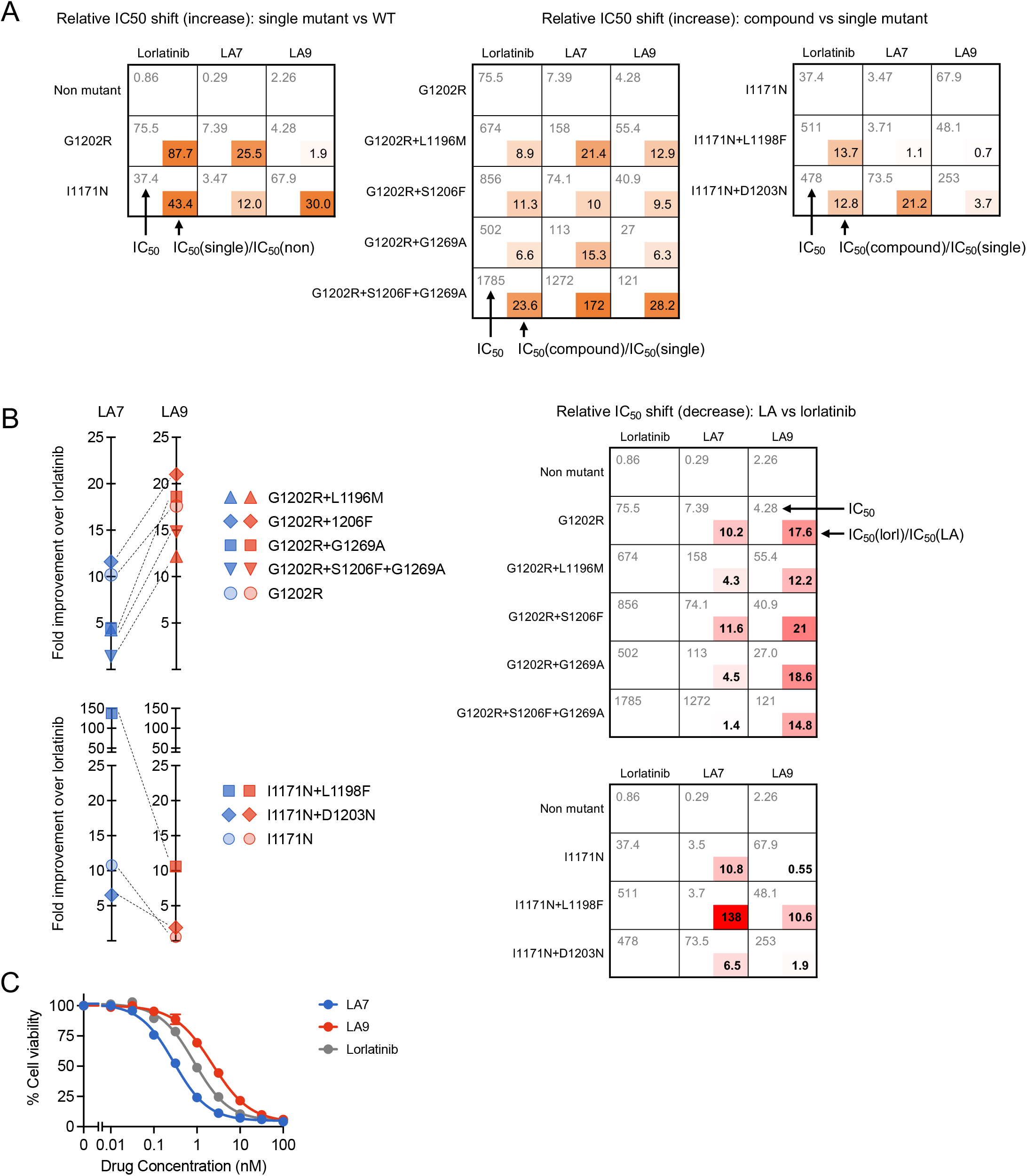
Relative changes in IC_50_ values for lorlatinib, LA7 and LA9 against single and compound ALK mutations. (A) Lorlatinib, LA7 and LA9 cellular IC_50_ values (upper left) and IC_50_ ratios (lower right) of single mutant vs nonmutant ALK (left panel) or compound mutant vs single mutant ALK (middle and right panels). IC_50_ values correspond to data shown in Figure S5. (B) Fold improvement of LA7 or LA9 compared to lorlatinib against single and compound mutations (calculated by dividing the lorlatinib IC_50_ by the LA IC_50_). The fold-improvement of LA7 and LA9 compared to lorlatinib against I1171N and G1202R, respectively (shown in light shaded symbols on left panels), is similar to that of compound mutations (darker shaded symbols). (C) Cell viability assays performed with Ba/F3 cells expressing nonmutant EML4-ALK with 0.01-100 nM lorlatinib, LA7 or LA9 for 48 hours. The viabilities were measured with CellTiter-Glo assay. Error bars indicate SE.

**Figure S14.**
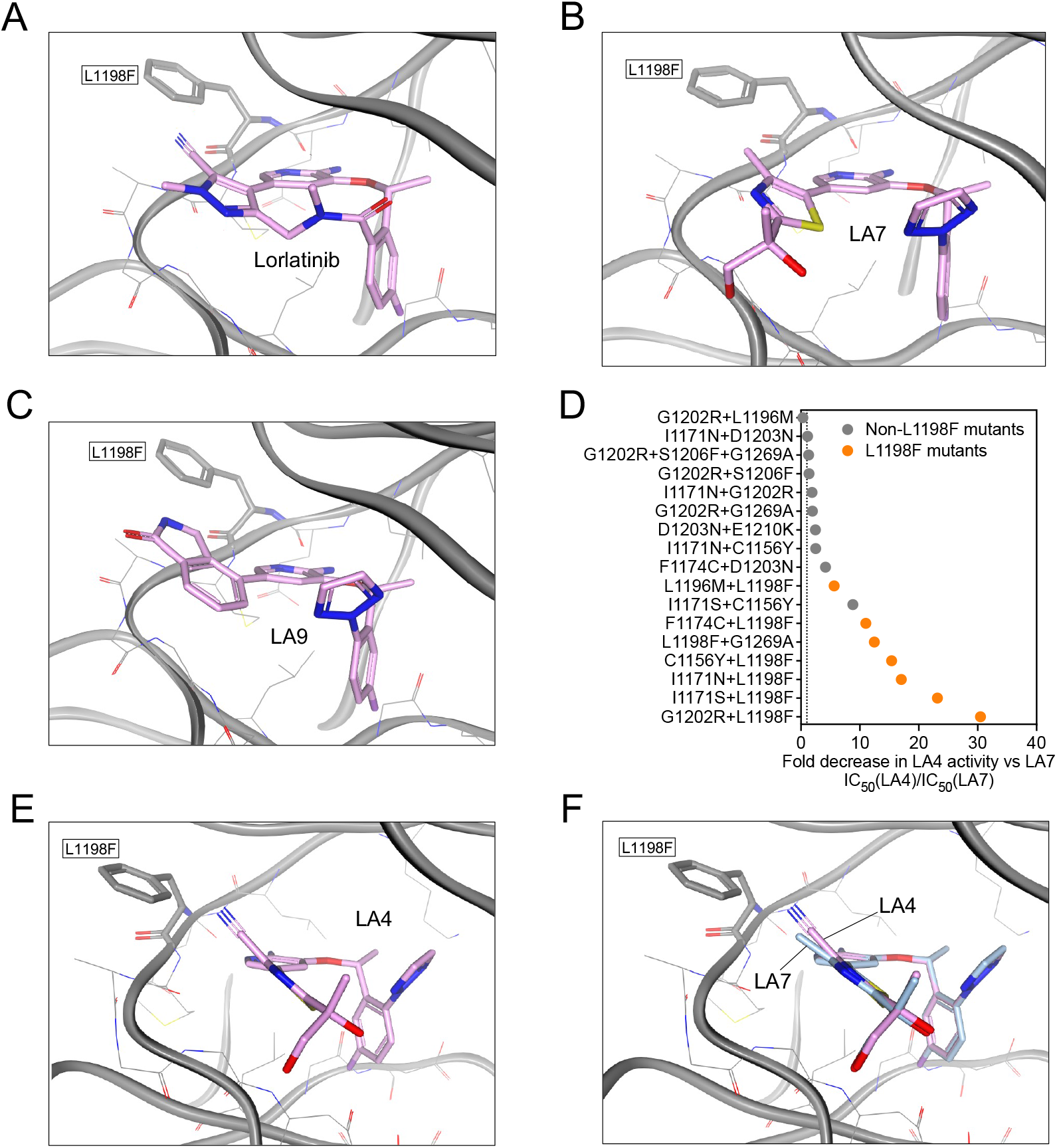
Structural considerations of the L1198F mutation. The L1198F mutation was modeled onto co-crystal structures of **l**orlatinib (A), LA7 (B), LA9 (C) or LA4 (E) bound to WT ALK. The L1198F substitution results in steric clash with the selectivity nitrile of lorlatinib. LA7 and LA9 lack the selectivity nitrile and can easily accommodate the L1198F substitution, whereas the nitrile of LA4 is positioned toward L1198F but exhibits reduced steric clash compared to lorlatinib due to the more flexible ligand structure. (D) Relative fold potency decrease of LA4 compared with LA7 (calculated by dividing the cellular IC_50_ values of LA4 by the that of LA7) against Ba/F3 models harboring compound ALK mutants. IC_50_ values correspond to data shown in Figure S5. (F) Superimposition of LA7 onto the LA4/L1198F model shown in Panel E comparing the position of the corresponding thiazole methyl or nitrile groups near L1198F.

